# Identification and Optimization of 4-Anilinoquinolines as Inhibitors of Cyclin G Associated Kinase

**DOI:** 10.1101/117630

**Authors:** Christopher R. M. Asquith, Tuomo Laitinen, James M. Bennett, Paulo H. Godoi, Graham J. Tizzard, Jonathan M. Elkins, Timothy M. Willson, William J. Zuercher

## Abstract

4-Anilinoquinolines were identified as potent and narrow spectrum inhibitors of the cyclin G associated kinase (GAK), an important regulator of viral and bacterial entry into host cells. Optimization of the 4-anilino group and the 6,7-quinoline substituents produced GAK inhibitors with nanomolar activity and over 50,000-fold selectivity relative to other members of the numb-associated kinase (NAK) sub-family. These compounds may be useful tools to explore the therapeutic potential of GAK in prevention of a broad range of infectious diseases.

## INTRODUCTION

Cyclin G associated kinase (GAK) is a ubiquitously expressed 160 kDa serine/threonine kinase involved in membrane trafficking. (Kanaoka, Kimura et al. 1997, Kimura, Tsuruga et al. 1997, Greener, Zhao et al. 2000, Korolchuk and Banting 2002). GAK was initially identified as a protein that associated with cell cycle regulator cyclin G (Kanaoka, Kimura et al. 1997). GAK knockdown by siRNA initiated cell-cycle arrest at the metaphase, demonstrating the requirement of GAK for normal mitotic progression (Shimizu, Nagamori et al. 2009). GAK is a member of the numb-associated kinase (NAK) family, which includes AAK1 (adaptor-associated kinase), STK16/MPSK1 (serine/threonine kinase 16/myristoylated and palmitoylated serine/threonine kinase 1), and BMP2K/BIKE (BMP-2 inducible kinase) (Manning, Whyte et al. 2002).

GAK and AAK1 are host cell kinases that regulate clathrin-mediated endocytosis, a critical regulatory process by which oligomeric clathrin and adaptor protein complexes facilitate entry of macromolecules, proteins and nutrients into cells (Barbieri, Di Fiore et al. 2016). GAK and AAK1 recruit clathrin and adaptor protein complexes to the cell membrane in part through phosphorylation of T156 in the μ2 subunit of adaptor protein-2 (AP-2). The clathrin/AP-2 complex facilitates vesicle assembly and efficient internalization of cargo proteins. In addition, GAK also regulates recycling of clathrin back to the cell surface, while AAK1 mediates the rapid recycling of receptors back to the plasma membrane. Through their involvement in clathrin-mediated endocytosis, GAK and AAK1 regulate EGFR internalization, thereby promoting EGF uptake and EGFR signaling (Neveu, Ziv-Av et al. 2015).

Clathrin-mediated endocytosis is a common mechanism by which viruses, toxins, and bacteria enter their host cells (Mercer, Schelhaas et al. 2010). Several viruses that add significant global disease burden, such as influenza, HCV, dengue, Hantaan virus, and Junin arenavirus, use this mechanism to infect cells (Mercer, Schelhaas et al. 2010). HIV, Ebola and Zika have also been characterized as entering the cell *via* this uptake pathway (Aleksandrowicz, Marzi et al. 2011, Sloan, Kuhl et al. 2013, Sikka, Chattu et al. 2016). In addition, the endocytosis of anthrax and diphtheria toxins is mediated by clathrin (Abrami, Liu et al. 2003, Papatheodorou, Zamboglou et al. 2010). Inhibition of NAK family kinases has been proposed as a new approach to development of host-centered anti-viral drugs (Bekerman and Einav 2015, Kovackova, Chang et al. 2015). A notable advantage of this approach is that these host molecular targets may be less prone to viral and bacterial resistance mechanisms.

GAK participates in a variety of other important biological processes through its roles in cell proliferation and receptor trafficking. GAK is a prognostic marker in advanced diseases, including hormone refractory prostate cancer (Ray, Wafa et al. 2006). GAK interacts directly with the androgen receptor (AR) and sensitizes it to low levels of androgens; the expression of GAK increases upon prolonged androgen treatment and during the progression of cells to hormone independence (Ray, Wafa et al. 2006, Susa, Choy et al. 2010, Sakurai, Ozaki et al. 2014). GAK is overexpressed in osteosarcoma cell lines and tissues where it contributes to proliferation and survival (Susa, Choy et al. 2010). Genome-wide association studies (GWAS) have identified single nucleotide polymorphisms in the GAK gene associated with susceptibility to Parkinson’s disease (Rhodes, Sinsheimer et al. 2011, Dzamko, Zhou et al. 2014). Additional studies support a functional role for GAK in the pathology of Parkinson’s disease—for example, enhanced toxicity was observed upon siRNA knockdown of GAK in HEK293 cells overexpressing α-synuclein, a major constituent of Lewy bodies, the protein aggregates in nerve cells of patients with Parkinson’s disease and other forms of dementia (Dumitriu, Pacheco et al. 2011, Dumitriu, Latourelle et al. 2012).

Prior reports have disclosed potent GAK inhibitors that lack selectivity over other protein kinases (Figure 1), such as the pyridinylimidazole p38 inhibitor SB203580, the c-Jun *N-*terminal kinase inhibitor SP600125, and dasatinib (Bellei, Pitisci et al. 2014). Intriguingly, the clinically-approved EGFR inhibitors gefitinib, erlotinib, and pelitinib show off target activity on GAK, with potency in the low nanomolar range (Fabian, Biggs et al. 2005). It is not known whether the clinical efficacy or adverse events observed with these kinase inhibitor drugs are connected to their inhibition of GAK activity. These drugs were designed as inhibitors of EGFR, and their utility as tools to study the biology of GAK is severely limited. Notably, it has been proposed that GAK inhibition causes clinical toxicity due to pulmonary alveolar dysfunction, but this controversial hypothesis has not been addressed with a selective small molecule GAK inhibitor (Tabara, Naito et al. 2011, Takada and Matsuo 2012).

**Figure 1.**
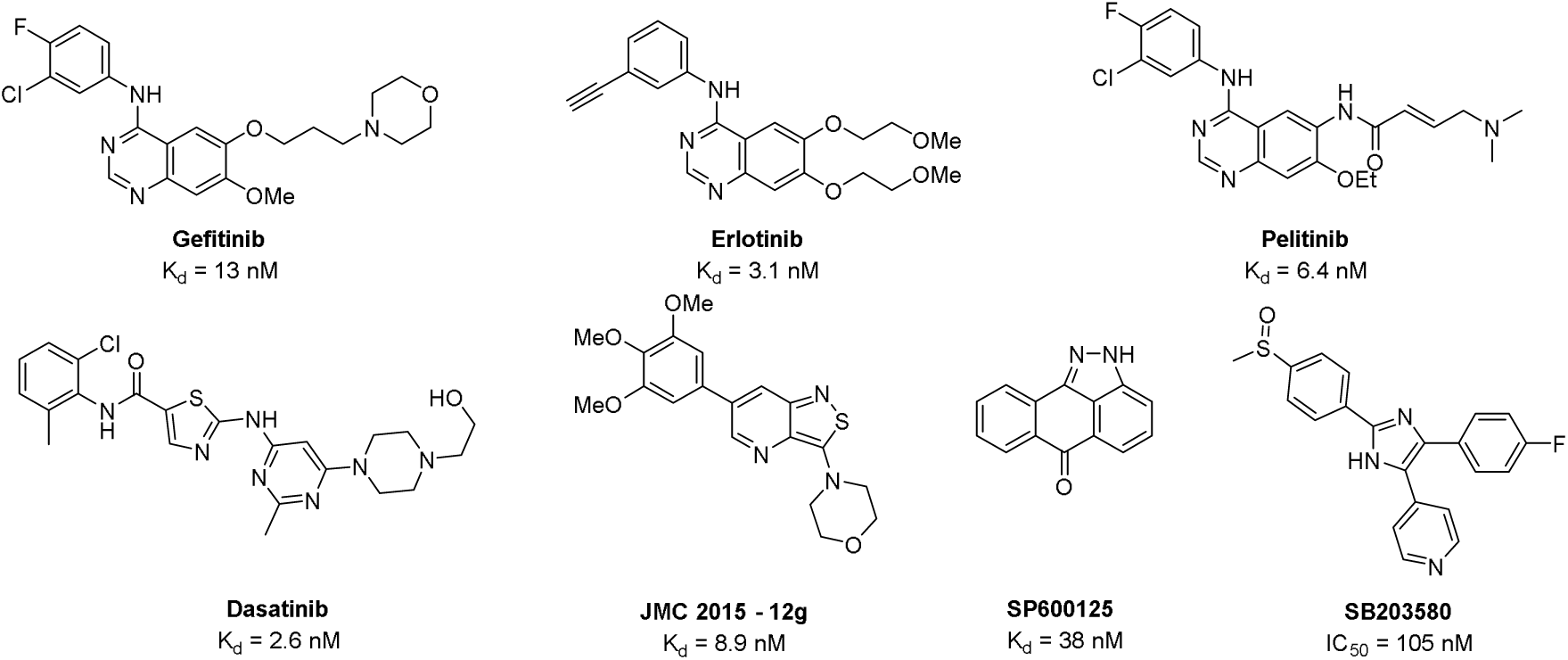
Known compounds with reported GAK activity

Recently isothiazolo[5,4-*b*]pyridines have been described as first generation chemical probes for GAK (Kovackova, Chang et al. 2015). Analog 12g was identified as a potent narrow spectrum GAK inhibitor with only a limited number of kinase off-targets including KIT, PDGFRB, FLT3, and MEK5. The availability of chemical probes with improved selectivity for GAK or a different spectrum of off-targets would be useful in target validation studies. Here we report the synthesis and characterization of 4-anilinoquinolines and 4-anilinoquinazolines as potent narrow spectrum GAK inhibitors. Several of these compounds have potential for development into high quality chemical probes for the study of GAK biology.

## RESULTS

The chemogenomic sets PKIS and PKIS2 contain hundreds of biologically and chemically diverse kinase inhibitors with potent activity across the protein kinome (Drewry, Wells et al. 2017). Through the screening of PKIS2 we identified several 4-anilinoquinolines and 4-anilinoquinazolines with nanomolar activity on GAK and varying degrees of selectivity with respect to other kinases (Table 1 and Figure 2). 4-Anilinoquinoline **1** had a K_d_ = 5.3 nM against GAK and showed potent activity on only one other kinase, ADCK3 (K_d_ = 220 nM). The analog **2**, containing a 4-pyridyl substituent at the 7-positon of the quinoline, retained GAK activity but also inhibited eighteen other kinases. The 4-anilinoquinazoline **3** showed a K_d_ = 15 nM on GAK with activity on only three additional protein kinases, EGFR (K_d_ = 0.32 nM), ERBB2 (K_d_ = 85 nM), and BUB1 (K_d_ = 130 nM). Our interest in the quinoline/quinazoline scaffold was heightened by the reported GAK activity of gefitinib, erlotinib, and pelitinib (Figure 1).

**Table 1.** Chemical starting points identified by screening PKIS2 Screening.

| Cmpd | X | R<sup>1</sup> | R<sup>2</sup> | R<sup>3</sup> | R<sup>4</sup> | R<sup>5</sup> | GAK<br>K<sub>d</sub> (nM) | Kinase off<br>targets<sup>a</sup> |
| --- | --- | --- | --- | --- | --- | --- | --- | --- |
| 1 | CH | OMe | OMe | OMe | CF<sub>3</sub> | H | 5.3 | 1 |
| 2 | CH | OMe | OMe | OMe | H | 4-pyridyl | 1.5 | 18 |
| 3 | N | Br | H | H | OMe | OMe | 15 | 3 |

<sup>a</sup>Number of kinases with over 95% binding at 1 μM in a Discoverx KINOMEScan experiment.

**Figure 2.**
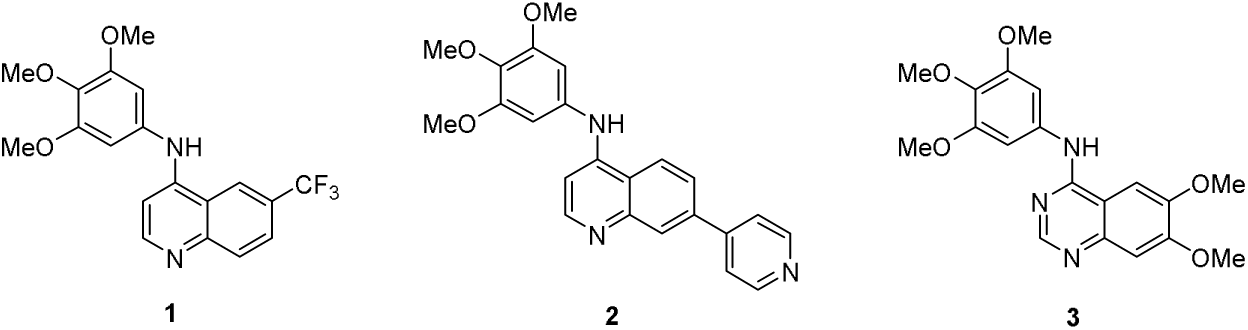
PKIS/PKIS2 quinoline and quinazoline hit compounds

To further explore the structural requirements for GAK activity, we synthesized analogs of **1** and **3** as outlined in Scheme 1. 4-Anilinoquinolines were prepared by heating the corresponding 4-chloroquinoline and substituted aniline in ethanol at reflux for 18 hours. Products were isolated by direct crystallization from the crude reaction mixture or following chromatographic purification. Most of the 4-anilinoquinazolines were prepared by heating the corresponding 4-chloroquinazoline and substituted aniline in *n*-butanol at reflux for 18 hours. The lower reactivity of the 4-chloroquinazoline required heating at the higher temperature, and these reactions were generally not as clean, requiring chromatographic purification to isolate the products. Several of the less nucleophilic anilines required an alternative method to be employed in which the aniline was added through a Buchwald-Hartwig coupling reaction to produce the corresponding 4-anilinoquinazolines **14** and **15** (Louie and Hartwig 1995).

**Scheme 1.**
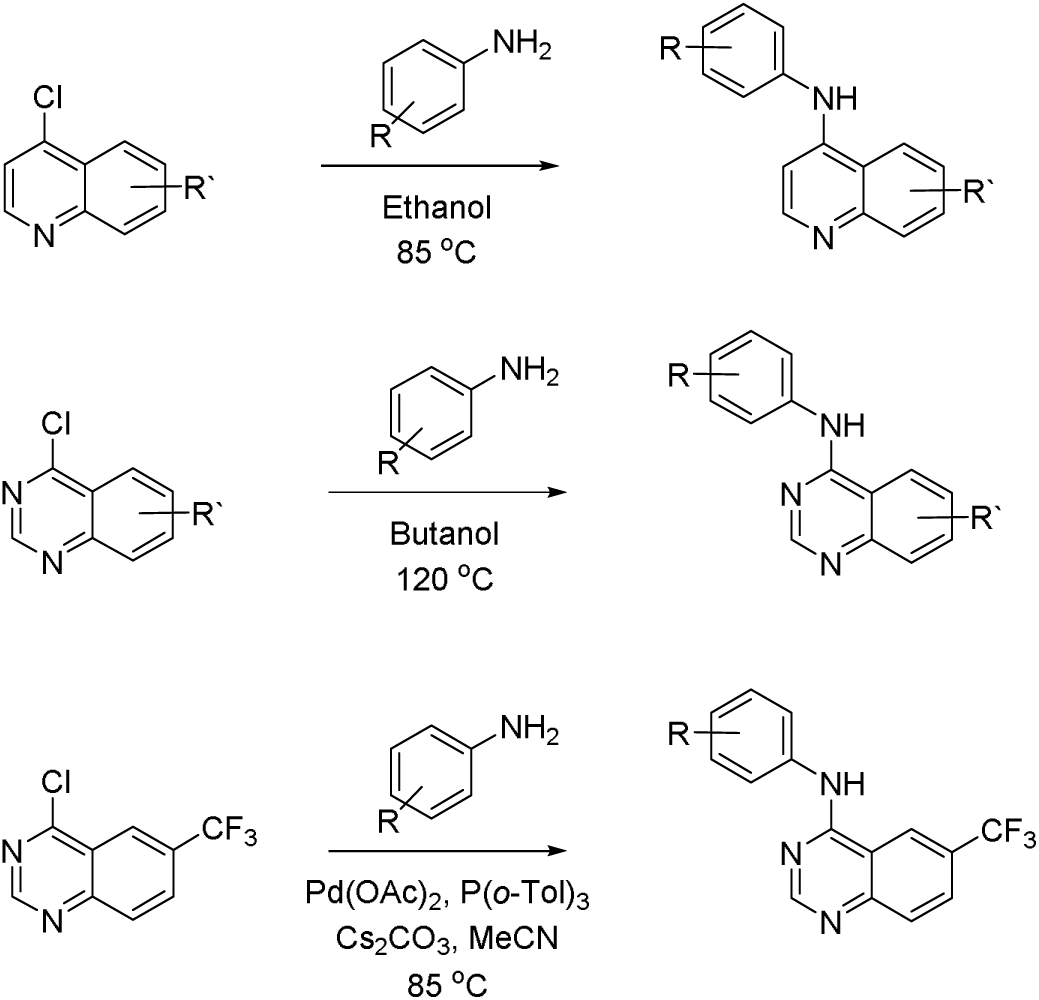
Preparation of quinoline and quinazoline based GAK inhibitors

The final compounds were initially screened at 10 μM for their ability to affect the melting temperature of the GAK protein using a differential scanning fluorimetry (DSF) assay. The change in denaturation temperature (ΔT_m_) in the presence and absence of a small molecule inhibitor was previously established as predictive of compound affinity (Fedorov, Niesen et al. 2012). The compounds were next screened for activity on the kinase domains of all four members of the NAK sub-family (GAK, AAK1, BMP2K, and STK16) using a TR-FRET binding displacement assay in a 16-point dose response format to determine the inhibition constant (K_i_).

The 4-anilinoquinoline **1** had ΔT_m_= 6.3 °C in the GAK DSF assay (Table 2). In the GAK binding displacement assay **1** demonstrated K_i_ = 3.9 nM with only weak activity on other members of the NAK family and a selectivity index, defined as the ratio of GAK K_i_ and the K_i_ of nearest NAK family member, over 4000 (Table 2). The first series of analogs explored deletion of one or more of the methoxy substituents. Removal of either the 4-methoxy (**4**) or 3-methoxy (**5**) group led to a 10-fold drop in potency against GAK and a slight increase in activity on other NAK family members. Modulating the position of the methoxy groups to 2,4-dimethoxy (**6**) or 2,5-dimethoxy (**7**) positions led to a decrease in GAK activity. The effect of substituting the aniline with a single methoxy group were more dramatic. Incorporating a single 4- methoxy group, as in **8**, resulted in a 100-fold drop in activity. However, analog **9** with a single 3-methoxy group retained potent GAK activity, demonstrating K_i_ = 5.7 nM and a NAK family selectivity index >2500. The 2-methoxy substituted **10** was an order of magnitude less potent than the parent compound **1** and maintained moderate NAK family selectivity. Two analogs were prepared in which the alkoxy substituents were bridged by a methylene (**11**) and an ethylene unit (**12).** However, both compounds showed a decrease in GAK activity compared to the corresponding 3,4-dimethoxy analog **6.**

**Table 2.** Small modifications to the hit compound - **1**

| Cmpd | X | R<sup>1</sup> | R<sup>2</sup> | R<sup>3</sup> | R<sup>4</sup> | GAK DSF<br>$\Delta T_m$ (°C)<sup>a</sup> | Binding displacement assay K<sub>i</sub> (μM) | | | | NAK<br>selectivity<sup>b</sup> |
| --- | --- | --- | --- | --- | --- | --- | --- | --- | --- | --- | --- |
|  |  |  |  |  |  |  | GAK | AAK1 | BMP2K | STK16 |  |
| **1** | CH | H | OMe | OMe | OMe | 6.3 | 0.0039 | 54 | >100 | 17 | 4300 |
| **4** | CH | H | OMe | H | OMe | 4.3 | 0.054 | 34 | 85 | 52 | 640 |
| **5** | CH | H | OMe | OMe | H | 3.4 | 0.058 | 24 | 44 | 24 | 420 |
| **6** | CH | OMe | H | OMe | H | 1.8 | 0.86 | 71 | >100 | 70 | 81 |
| **7** | CH | OMe | H | H | OMe | 2.6 | 0.14 | 80 | >100 | 58 | 420 |
| **8** | CH | H | H | OMe | H | 1.7 | 0.73 | 56 | 75 | 19 | 26 |
| **9** | CH | H | OMe | H | H | 5.8 | 0.0057 | 14 | 23 | 18 | 2500 |
| **10** | CH | OMe | H | H | H | 3.0 | 0.044 | 4.8 | 11 | 29 | 110 |
| **11** | CH | H | OCH<sub>2</sub>O |  | H | 3.2 | 0.29 | 30 | 60 | 25 | 85 |
| **12** | CH | H | OCH<sub>2</sub>CH<sub>2</sub>O |  | H | 1.5 | 0.88 | 44 | 56 | 13 | 14 |
| **13** | N | H | OMe | OMe | OMe | 2.4 | 0.037 | 7.8 | 18 | 2.7 | 73 |
| **14** | N | H | OMe | H | OMe | 0.7 | 0.31 | >100 | >100 | 8.0 | 26 |
| **15** | N | H | H | OMe | H | 0.3 | 1.8 | 11 | >100 | 2.7 | 1.5 |

<sup>a</sup>Average of 4 experiments
<sup>b</sup>Selectivity against the nearest NAK family member

Switching to the 4-anilinoquinazoline core, the effect of variation in methoxy substitution was explored in three analogs (Table 1). The rank order activity of the mono-, di-, and trimethoxy 4-anilinoquinazolines **13**–**15** paralleled that of the corresponding 4- anilinoquinoline analogs **1, 4**, and **5**, but the quinazolines showed a 10-fold decrease in potency compared with the quinolines. The 3,4,5-trimethoxyanilino analog **13** was the most potent on GAK but showed a greatly reduced NAK family selectivity. The lower potency and selectivity of the 4-anilinoquinazoline core led us to focus our efforts on the 4-anilinoquinolines.

The next series of 4-anilinoquinoline analogs explored the variation in electronic properties and steric bulk of the aniline substituents (Table 3). Replacing the methoxy groups with fluorine yielded several potent compounds. Mono-, di-, and trifluoro analogs **16**-**20** showed potent activity on GAK but with reduced NAK family selectivity. Analogs with chlorine substitution (**21**–**24**) also retained potent GAK activity with the exception of the 2-chloro derivative **24**. The bromo- and iodo-substituted analogs **27**–**30** were less potent. Non-halogen electron withdrawing substituents were generally poorly tolerated: cyano (**31**–**33**), trifluromethyl (**34**), and methylsulphonyl (**37**–**38**) analogs showed reduced activity. The bulky 4-*tert*-butoxy and 4-methylsulphonylmethyl analogs **39** and **40** had weaker activity. The only remaining compound with potent GAK activity was the 3-acetyl analog **36**, but it showed modest NAK family selectivity.

**Table 3.**
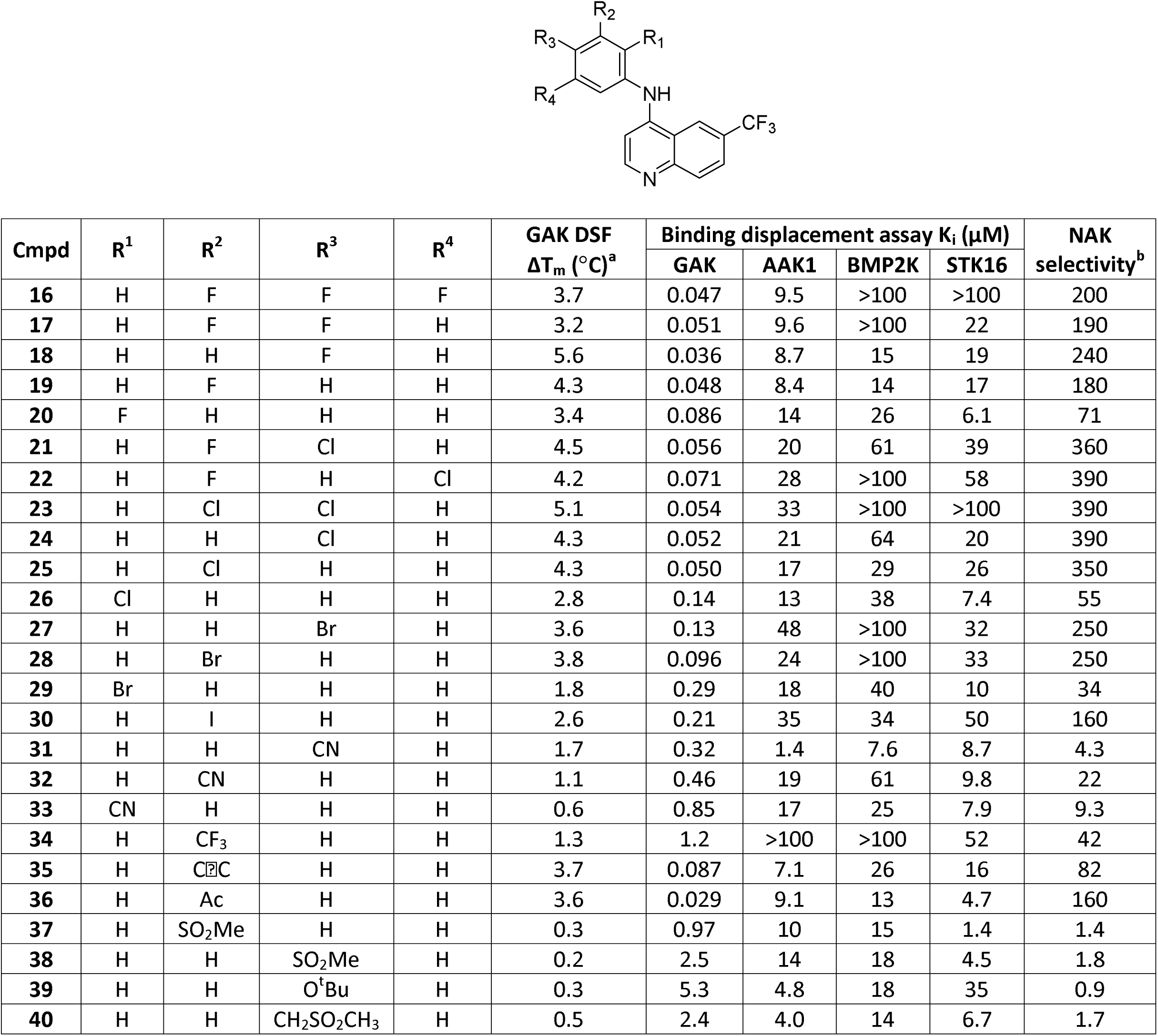
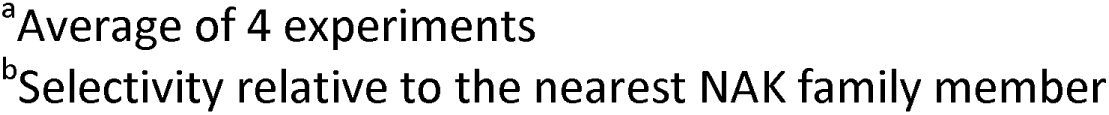
Modifications to the aniline segment of compound - **1**

| Cmpd | R<sup>1</sup> | R<sup>2</sup> | R<sup>3</sup> | R<sup>4</sup> | GAK DSF<br>ΔT<sub>m</sub> (°C)<sup>a</sup> | Binding displacement assay K<sub>i</sub> (μM) |  |  |  | NAK<br>selectivity<sup>b</sup> |
| --- | --- | --- | --- | --- | --- | --- | --- | --- | --- | --- |
|  |  |  |  |  |  | GAK | AAK1 | BMP2K | STK16 |  |
| 16 | H | F | F | F | 3.7 | 0.047 | 9.5 | >100 | >100 | 200 |
| 17 | H | F | F | H | 3.2 | 0.051 | 9.6 | >100 | 22 | 190 |
| 18 | H | H | F | H | 5.6 | 0.036 | 8.7 | 15 | 19 | 240 |
| 19 | H | F | H | H | 4.3 | 0.048 | 8.4 | 14 | 17 | 180 |
| 20 | F | H | H | H | 3.4 | 0.086 | 14 | 26 | 6.1 | 71 |
| 21 | H | F | Cl | H | 4.5 | 0.056 | 20 | 61 | 39 | 360 |
| 22 | H | F | H | Cl | 4.2 | 0.071 | 28 | >100 | 58 | 390 |
| 23 | H | Cl | Cl | H | 5.1 | 0.054 | 33 | >100 | >100 | 390 |
| 24 | H | H | Cl | H | 4.3 | 0.052 | 21 | 64 | 20 | 390 |
| 25 | H | Cl | H | H | 4.3 | 0.050 | 17 | 29 | 26 | 350 |
| 26 | Cl | H | H | H | 2.8 | 0.14 | 13 | 38 | 7.4 | 55 |
| 27 | H | H | Br | H | 3.6 | 0.13 | 48 | >100 | 32 | 250 |
| 28 | H | Br | H | H | 3.8 | 0.096 | 24 | >100 | 33 | 250 |
| 29 | Br | H | H | H | 1.8 | 0.29 | 18 | 40 | 10 | 34 |
| 30 | H | I | H | H | 2.6 | 0.21 | 35 | 34 | 50 | 160 |
| 31 | H | H | CN | H | 1.7 | 0.32 | 1.4 | 7.6 | 8.7 | 4.3 |
| 32 | H | CN | H | H | 1.1 | 0.46 | 19 | 61 | 9.8 | 22 |
| 33 | CN | H | H | H | 0.6 | 0.85 | 17 | 25 | 7.9 | 9.3 |
| 34 | H | CF<sub>3</sub> | H | H | 1.3 | 1.2 | >100 | >100 | 52 | 42 |
| 35 | H | C≡C | H | H | 3.7 | 0.087 | 7.1 | 26 | 16 | 82 |
| 36 | H | Ac | H | H | 3.6 | 0.029 | 9.1 | 13 | 4.7 | 160 |
| 37 | H | SO<sub>2</sub>Me | H | H | 0.3 | 0.97 | 10 | 15 | 1.4 | 1.4 |
| 38 | H | H | SO<sub>2</sub>Me | H | 0.2 | 2.5 | 14 | 18 | 4.5 | 1.8 |
| 39 | H | H | O<sup>t</sup>Bu | H | 0.3 | 5.3 | 4.8 | 18 | 35 | 0.9 |
| 40 | H | H | CH<sub>2</sub>SO<sub>2</sub>CH<sub>3</sub> | H | 0.5 | 2.4 | 4.0 | 14 | 6.7 | 1.7 |

<sup>a</sup>Average of 4 experiments
<sup>b</sup>Selectivity relative to the nearest NAK family member

Having explored the steric and electronic requirements of the aniline group, our focus turned to the quinoline core (Table 4). The effect of varying the 6- and 7- substituent on the quinoline core was explored within the 3,4,5-trimethoxyaniline series. Removal of the trifluromethyl group (**41**) resulted in a 10-fold drop in NAK family potency, although selectivity for GAK was retained. The 6-fluoro (**42**) and 7-fluoro (**43**) substituted analogs demonstrated potent GAK activity and >10,000-fold NAK family selectivity. In contrast, the 5,7-difluoro compound **44** showed a significant drop in GAK activity compared with the 7- fluoro compound **43**. Compounds with 6-tert-butyl (**45**), 6-cyano (**46**), and 6-sulfonylmethyl (**47**) substituents demonstrated potent GAK and NAK family selectivity >1000-fold. While 6- methoxy substitution (**48**) resulted in a slight decrease in GAK activity, 6,7-dimethoxy substitution (**49**) yielded a dramatic increase in GAK activity and NAK family selectivity. Compound **49** showed >8 °C stabilization of GAK protein in the thermal denaturation assay and K_i_ = 0.53 nM in the TR-FRET assay with NAK family selectivity >50,000. Finally, the 7- substituted analogs **50**–**52** showed single digit nanomolar GAK potency and NAK family selectivity of approximately 10,000-fold.

**Table 4.** Modifications to the quinoline core of compound - **1**

| Cmpd | R<sup>1</sup> | R<sup>2</sup> | R<sup>3</sup> | GAK DSF<br>$\Delta T_m$ (°C)<sup>a</sup> | Binding displacement assay K<sub>i</sub> (μM) | | | | NAK<br>selectivity<sup>b</sup> |
| --- | --- | --- | --- | --- | --- | --- | --- | --- | --- |
|  |  |  |  |  | GAK | AAK1 | BMP2K | STK16 |  |
| **1** | H | CF<sub>3</sub> | H | 6.3 | 0.0039 | 54 | >100 | 17 | 4300 |
| **41** | H | H | H | 4.4 | 0.040 | >100 | >100 | >100 | >2500 |
| **42** | H | F | H | 5.9 | 0.0057 | 83 | >100 | >100 | 14000 |
| **43** | H | H | F | 5.8 | 0.0024 | 44 | >100 | >100 | 18000 |
| **44** | F | H | F | 3.1 | 0.14 | >100 | >100 | >100 | >720 |
| **45** | H | <sup>t</sup>Bu | H | 5.2 | 0.0081 | >100 | >100 | 68 | 8400 |
| **46** | H | CN | H | 4.4 | 0.0015 | 16 | 16 | 2.5 | 11000 |
| **47** | H | SO<sub>2</sub>Me | H | 3.3 | 0.018 | 73 | >100 | 61 | 3400 |
| **48** | H | OMe | H | 6.0 | 0.013 | >100 | >100 | >100 | >7700 |
| **49** | H | OMe | OMe | 8.8 | 0.00054 | 28 | 63 | >100 | 51000 |
| **50** | H | H | OMe | 6.1 | 0.0026 | 58 | >100 | >100 | 22000 |
| **51** | H | H | CF<sub>3</sub> | 5.6 | 0.0062 | 90 | >100 | >100 | 15000 |
| **52** | H | H | CN | 5.3 | 0.0027 | 67 | >100 | 25 | 9300 |

<sup>a</sup>Average of 4 experiments<sup>b</sup>Selectivity relative to the nearest NAK family member

A final series of analogs explored the effect of matching the potent 6,7-dimethoxy core with a range of 4-anilino substituents (Table 5). The 3-methoxy analog **9** retained the potency of the trimethoxy analog **1** in the original series (Table 2). When 3-methoxyaniline was incorporated into the 6,7-dimethoxyquinoline core (**53**), there was a 4-fold reduction in GAK activity compared to the trimethoxy analog **49**. The 3-bromo (**54**) and 3-acetylene (**55**) substituted 4-anilino-6,7-dimethoxyquinolines showed similar drops in GAK potency and decreases in NAK family selectivity. The 3-acetyl substituent (**56**) led to a 100-fold loss of GAK activity relative to **49**, and this compound was equipotent across the NAK family. Finally, the 6,7-dimethoxyquinazoline **58**, incorporating a 3,4,5-trimethoxyaniline, showed a two-fold loss in potency while maintaining a high selectivity relative to other NAK family members.

**Table 5.**
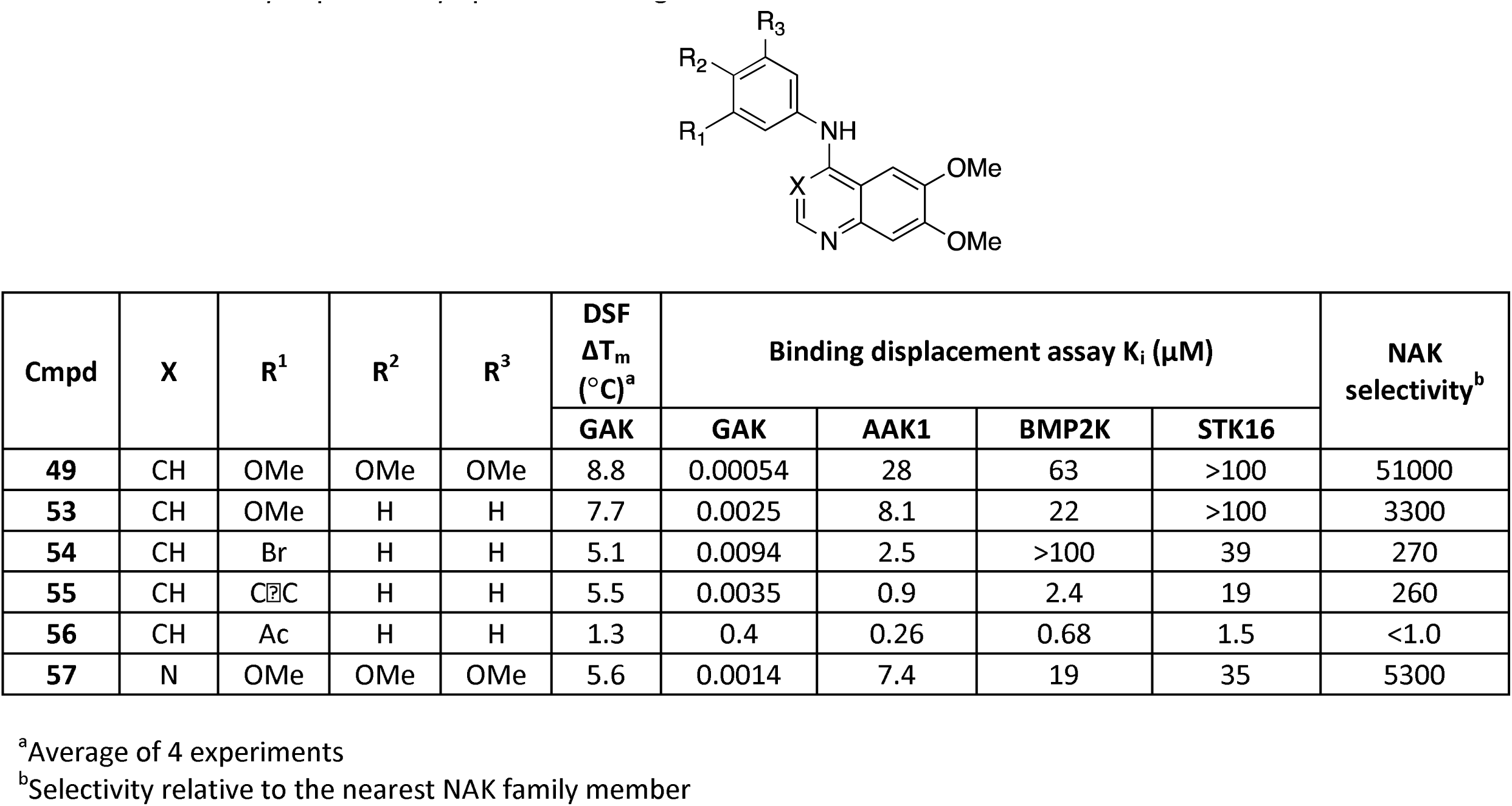
Small library of potentially optimized analogs of **55**

| Cmpd | X | R<sup>1</sup> | R<sup>2</sup> | R<sup>3</sup> | DSF<br>ΔT<sub>m</sub><br>(°C)<sup>a</sup> | Binding displacement assay K<sub>i</sub> (μM) |  |  |  | NAK<br>selectivity<sup>b</sup> |
| --- | --- | --- | --- | --- | --- | --- | --- | --- | --- | --- |
|  |  |  |  |  | GAK | GAK | AAK1 | BMP2K | STK16 |  |
| **49** | CH | OMe | OMe | OMe | 8.8 | 0.00054 | 28 | 63 | >100 | 51000 |
| **53** | CH | OMe | H | H | 7.7 | 0.0025 | 8.1 | 22 | >100 | 3300 |
| **54** | CH | Br | H | H | 5.1 | 0.0094 | 2.5 | >100 | 39 | 270 |
| **55** | CH | C≡C | H | H | 5.5 | 0.0035 | 0.9 | 2.4 | 19 | 260 |
| **56** | CH | Ac | H | H | 1.3 | 0.4 | 0.26 | 0.68 | 1.5 | <1.0 |
| **57** | N | OMe | OMe | OMe | 5.6 | 0.0014 | 7.4 | 19 | 35 | 5300 |

<sup>a</sup>Average of 4 experiments
<sup>b</sup>Selectivity relative to the nearest NAK family member

To rationalize the pivotal activity of the methoxy group on the aniline ring, we used molecular modelling to analyse the effect of inhibitor binding to the GAK catalytic site. Analogs **1, 5**, **9**, and **49** were docked into the active site of the GAK x-ray structure, and a hydration site analysis was performed with WaterMap using multiple orientations and conformations of each compound (Figure 3) (Young, Abel et al. 2007, Abel, Young et al. 2008, Kovackova, Chang et al. 2015). These calculations identified a network of water molecules that could occupy the active site. Docking of the 4-aminoquinolines into the site resulted in potential displacement of one specific water molecule that could contribute up to 6.4 kcal/mol in binding energy (Figure 3). The trimethoxy substituted anilines (**49** and **1**) were predicted to displace this water molecule most effectively (Figure 3A/3B and 3C respectively), whereas other methoxy substitution patterns such as the 3,4-dimethoxy (**5**) were only partially able to displace the water molecule (Figure 3D/3E). The water site analysis also explained why the meta-methoxy analog **9** was more potent than the ortho-and para- analogs **8** and **10. 9** could fully displace the high-energy water molecule (Figure 3F), whereas **8** and **10** were not able to do so.

**Figure 3.**
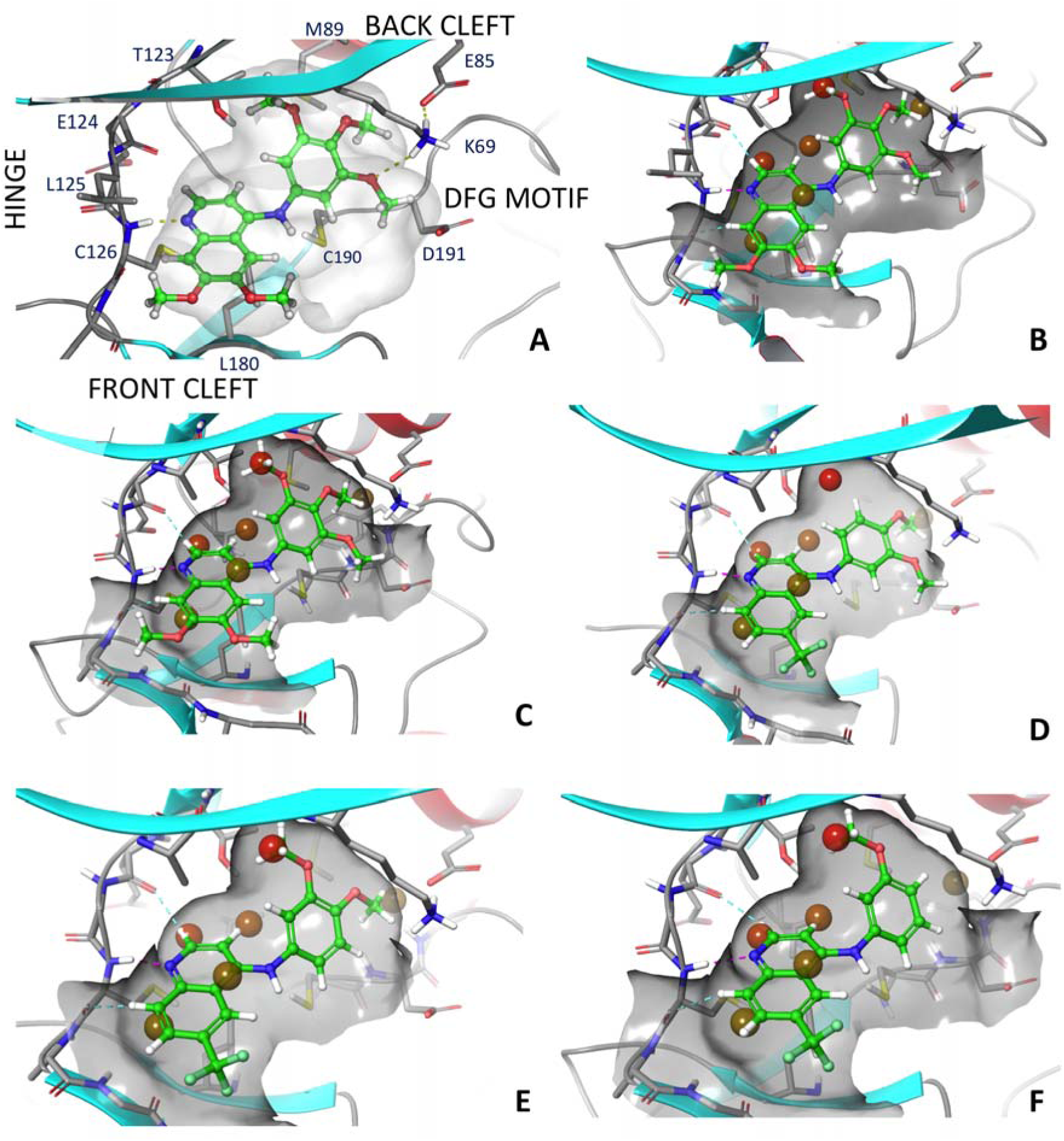
Active site WaterMap of compounds in GAK: A - Visualised GAK active site interaction of **49**, B - WaterMap of **49**, C - WaterMap of **1**, D/E - WaterMap of **5** favoured (D) and unfavoured (E), F - WaterMap of **9**.

We measured GAK affinity of the 4-anilinoquinoline analogs with a wide range of ΔT_m_ values and affinity over several orders of magnitude in the TR-FRET binding assay. The two binding assays were highly correlated, as shown by a plot of ΔT_m_ versus −log K_i_ which has R = 0.86 (Figure 4).

**Figure 4.**
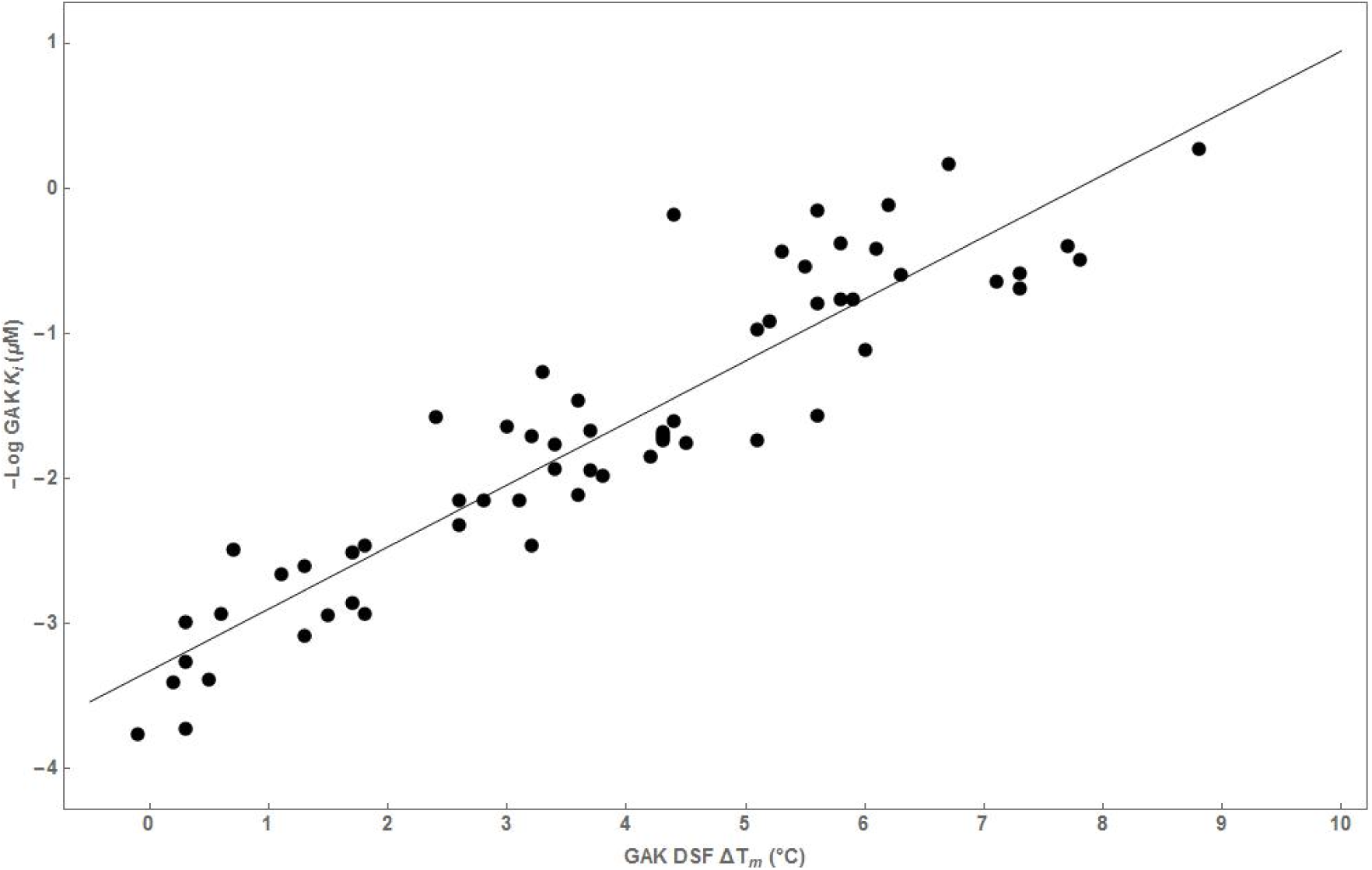
Correlation of GAK DSF ΔT_m_ vs -Log GAK K_i_ (μM)

## DISCUSSION

Chemical tractability is a hallmark of the protein kinases. Over 30 drugs have been approved that target the ATP cofactor binding site, predominantly in the field of oncology. However, most of these drugs were developed as multi-kinase inhibitors that leverage the conservation of the ATP pocket across the enzymes to increase their clinical efficacy. Development of kinase inhibitors for treatment of chronic diseases, such as inflammation and neurodegeneration, will require drugs with greatly improved selectivity profiles (Cohen and Alessi 2013). New chemical approaches and molecular insights into the development of highly selective kinase inhibitors that target the conserved ATP-binding site are urgently needed. Binding assays are the most accurate and robust method to measure potency and selectivity of ATP-competitive kinase inhibitors (Rudolf, Skovgaard et al. 2014). Moreover, binding affinity provides an accepted surrogate for direct measurement of enzyme inhibition in drug optimization studies. Binding assays available at commercial vendors are routinely used to profile kinase inhibitors for their selectivity across the human kinome (Davis, Hunt et al. 2011). Here, we have uncovered a series of 4-anilinoquinolines that show high affinity for GAK as measured in two orthogonal biochemical assays (DSF and TR-FRET), which were highly correlated (Figure 4). Some of these compounds demonstrated remarkable binding selectivity for GAK over hundreds of other kinases and even for other members of the closely related NAK sub-family. Using medicinal chemistry and molecular modelling, we have delineated the structural features of these molecules that contribute to the binding selectivity within the NAK sub-family.

Common structural elements of kinase ligand interactions have been systematically described in the KLIFS database (van Linden, Kooistra et al. 2014, Kooistra, Kanev et al. 2016). Briefly, kinase domains have *C*- and *N*-terminal domains bridged by a hinge region. ATP-competitive inhibitors generally make one to three hydrogen bonding interactions with the backbone residues in the hinge. The ATP binding site can be divided into a front cleft projecting towards solvent and a back cleft that has some degree of conformational plasticity, due in part to the disposition of a conserved loop containing an Asp-Phe-Gly (“DFG”) motif. The ease of access to the back cleft is largely determined by the size of the amino acid directly preceding the hinge, which is known as the “gatekeeper” residue.

Compounds containing quinoline and quinazoline ring systems have been the focus of prior kinase drug discovery campaigns, which in some cases have led to approved medicines (Bridges 1999, Solomon and Lee 2011, Ravez, Castillo-Aguilera et al. 2015). Structural studies using x-ray crystallography have been reported for several quinoline and quinazoline-based kinase inhibitors in complex with their target kinases (Wu, Nielsen et al. 2015). The dominant binding mode of these quinolines and quinazolines shows a hydrogen bond in the hinge region between N1 of the ligand and the third residue from the gatekeeper (GK + 3: Cys in GAK, BMP2K, and AAK1 and Phe in STK16) (Figure 3A). The aniline group at the 4-position of the quinazoline or quinoline ring system projects into the back-cleft pocket of GAK that is partially defined by the αC helix and the DFG motif. Finally, substituents at the 6- and 7-position of the quinoline are directed toward the surface of the protein. Changes in the aniline head group and on the quinoline core are known to affect kinase potency and selectivity profiles in other, structurally-related inhibitor series (Haile, Votta et al. 2016), as these parts of the ligand project into a pocket lined with varied functionality or having a range of conformational plasticity (van Linden, Kooistra et al. 2014).

We observed that GAK potency and NAK family selectivity was affected by a) the hinge interacting moiety (quinoline vs. quinazoline), b) the pendant aniline, and c) substituents at the quinoline 6- and 7-positions. The quinoline **1** was an order of magnitude more potent on GAK than the quinazoline **13**. The hydrogen that projects from C3 of the quinoline restricts the torsion angle defined with the aniline to a much greater degree than the quinazoline N3, from near planar for quinazoline up to 60° for quinoline. Quinoline 1 may be more conformationally restricted than **13**, a proposition supported by gas phase OPLS3 torsional scanning (Figure S1) and Density Function Theory (DFT) optimised (B3LYP/6-31G**) force field calculations (Table S1). These results are consistent with small molecule x-ray structures determined for **1, 9, 13, 17, 37, 48**, and **49** (Figure S2 and Table S2). Thus, the restricted conformation of the quinoline relative to the quinazoline may favourably preorganize **1** for binding to GAK (Figure 5).

**Figure 5.**
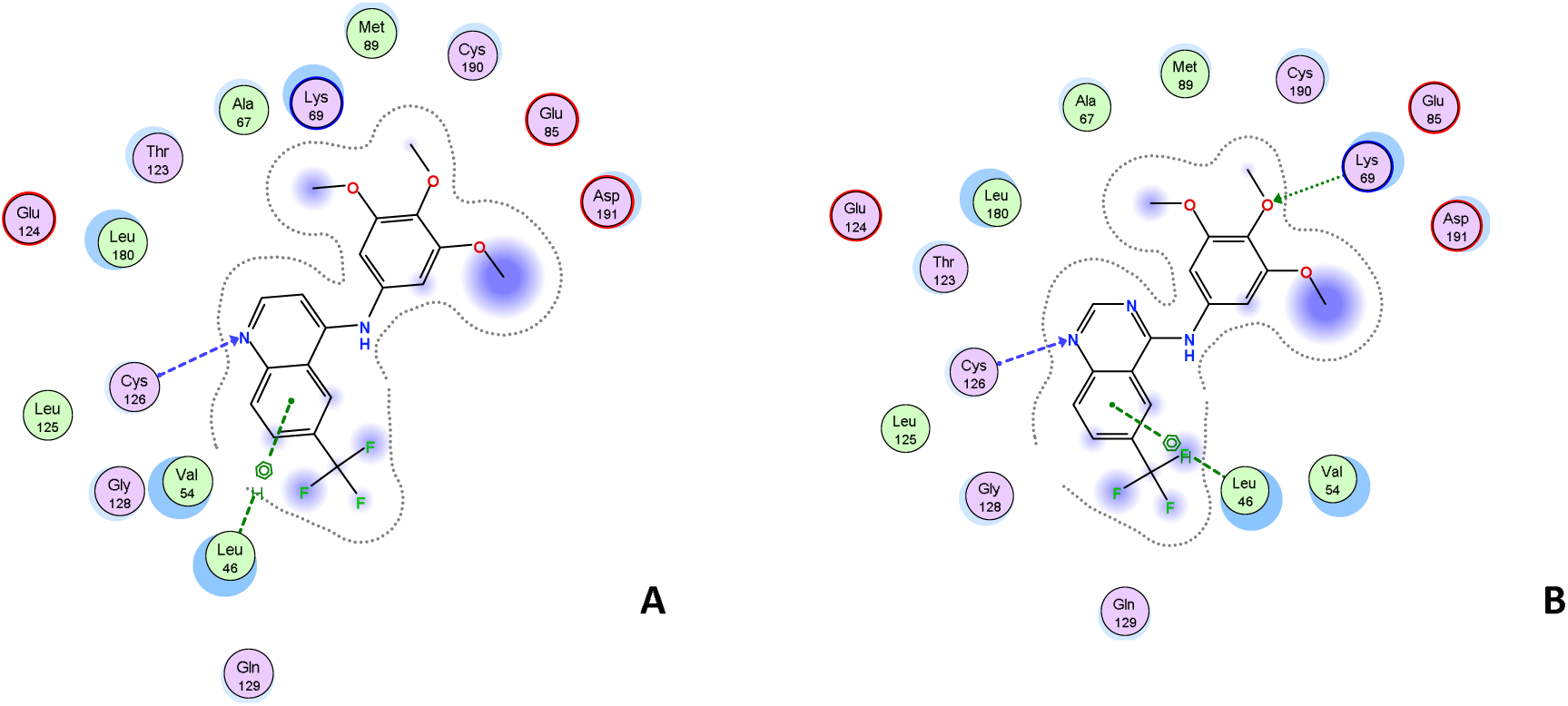
Comparison of GAK binding of compound **1** (A) and **13** (B). The quinoline scaffold is a tighter binder then the quinazoline despite the possibility for an additional hinge binding interaction.

WaterMap analysis of the binding site suggested that a coordinated water network in the protein pocket spans the region proximal the aniline and the 6-position of the ring system. We propose that compounds able to displace a poorly coordinated water molecule in this system will have increased potency due to entropic and enthalpic contributions to the free energy of binding (Figures 3 and S3). Compounds **1** and **16** are calculated to have their methoxy or fluoro substituents directly occupying the same space as the high-energy water molecule. On the other hand, **6** and **20** were comparatively weaker binders and are predicted to be unable to displace the water molecule. These models suggest that a water network within the GAK active site plays a critical role in defining the relative affinity of quinoline ligands. Extension of this model to other NAK family members or more distantly related kinases may lead to computational models to predict selectivity.

Variation of the substitution at the 6- and 7-positions was observed to have a large effect on GAK affinity. Comparing compounds in the 3,4,5,-trimethoxyaniline series, the compound with the weakest affinity was the unsubstituted quinoline **41**. The 6- and 7-positions are oriented towards solvent and may provide an opportunity to improve molecular properties. Indeed, the approved medicines gefitinib and erlotinib both incorporate water solubilizing groups at the 6- and 7-positions of their quinazoline cores (Figures S4-S7).

As a result of screening the four NAK family kinases in parallel, we also identified quinolines with interesting dual activity profiles (Figure S8). Most analogs had GAK affinity that was >100-fold higher than for the other NAK family kinases. However, there were some exceptions. **39**, which has a 4-*tert*-butoxy aniline substituent, had affinity for all four NAK family kinases within a range of 8-fold. Compound **37** with a 3-SO_2_Me aniline substituent had a notable potency boost on STK16, to the extent that its affinity was within 2-fold of GAK. These bulky substituents are likely to bind into the back cleft of the active site. In addition, it was also observed that a number of compounds including **36** and **56** have pan-NAK activity (Figure S9). Further exploration of this pocket may yield quinolines with selectivity for other members of the NAK family.

The measurement of binding affinity across the NAK family kinases in parallel allowed us to explore the determinants of selectivity within four close phylogenetic relatives. Inhibitors of the NAK family kinases have a high likelihood of showing low selectivity across the family based on high identity of their primary amino acid sequence and the presence of structural features which are distinct from those observed in other kinases. Specifically, the four NAK family kinases have a large α-helical insert positioned *C*-terminal to the activation segment, in proximity to the ATP binding site (Sorrell, Szklarz et al. 2016). Notably, GAK does have one fewer amino acid residue in the hinge region, which distinguishes it from the other NAK family kinases. Prior results with 144 clinically-used kinase inhibitors revealed few compounds selective for GAK across the NAK family. It is therefore remarkable that several of the 4-aminoquinolines bound to GAK with over 10,000-fold higher affinity compared to the other NAK family kinases.

In summary, the 4-anilinoquinoline **1** was identified as a narrow spectrum GAK inhibitor with remarkable selectivity within the NAK sub-family. In addition, broad screening in a commercial kinase panel found that ADCK3 was the only kinase within 30-fold affinity of GAK. Iterative medicinal chemistry led to a member of the 4-anilinoquinoline series with NAK family selectivity over 50,000-fold. Our results demonstrate that quinoline-based ATP-competitive kinase inhibitors can be designed with exquisite selectivity. More importantly, the 4-anilinoquinolines series exemplified by **1** has the potential to yield high quality chemical probes for use in the elucidation of GAK function in cells, and perhaps *in vivo*.

## EXPERIMENTAL

### General Procedures

All reactions were performed using flame-dried round-bottomed flasks or reaction vessels unless otherwise stated. Where appropriate, reactions were carried out under an inert atmosphere of nitrogen with dry solvents, unless otherwise stated. Dry acetonitrile (MeCN) was obtained directly from manufacturer and used without further purification. Yields refer to chromatographically and spectroscopically pure isolated yields. Reagents were purchased at the highest commercial quality and used without further purification, unless otherwise stated. Reactions were monitored by thin-layer chromatography carried out on 0.25 mm E. Merck silica gel plates (60F-254) using ultraviolet light as visualizing agent. NMR spectra were recorded on a Varian Inova 400 or Inova 500 spectrometer and were calibrated using residual undeuterated solvent as an internal reference (CDCl_3_: ^1^H NMR = 7.26, ^13^C NMR = 77.16). The following abbreviations or combinations thereof were used to explain the multiplicities observed: s = singlet, d = doublet, t = triplet, q = quartet, m = multiplet, br = broad. Liquid Chromatography (LC) and High Resolution Mass Spectra (HRMS) were recorded on a ThermoFisher hybrid LTQ FT (ICR 7T). The University of Southampton (Southampton, UK) small molecule X-ray facility collected and analyzed all X-ray diffraction data.

#### Mass Spectrometry

Samples were analyzed with a hybrid LTQ FT (ICR 7T) (ThermoFisher, Bremen, Germany) mass spectrometer coupled with a Waters Acquity H-class liquid chromatograph system. Samples were introduced *via* an electrospray source at a flow rate of 0.6 mL/min. Electrospray source conditions were set as: spray voltage 4.7 kV, sheath gas (nitrogen) 45 arb, auxiliary gas (nitrogen) 30 arb, sweep gas (nitrogen) 0 arb, capillary temperature 350°C, capillary voltage 40 V and tube lens voltage 100 V. The mass range was set to 150-2000 m/z. All measurements were recorded at a resolution setting of 100,000.

Separations were conducted on a Waters Acquity UPLC BEH C18 column (2.1 × 50 mm, 1.7 μM particle size). LC conditions were set at 100 % water with 0.1% formic acid (A) ramped linearly over 9.8 mins to 95% acetonitrile with 0.1% formic acid (B) and held until 10.2 mins. At 10.21 mins the gradient was switched back to 100% A and allowed to re-equilibrate until 11.25 mins.

Xcalibur (ThermoFisher, Breman, Germany) was used to analyze the data. Solutions were analyzed at 0.1 mg/mL or less based on responsiveness to the ESI mechanism. Molecular formula assignments were determined with Molecular Formula Calculator (v 1.2.3). Low-resolution mass spectrometry (linear ion trap) provided independent verification of molecular weight distributions. All observed species were singly charged, as verified by unit m/z separation between mass spectral peaks corresponding to the ^12^C and ^13^ C^12^ C_c-1_ isotope for each elemental composition.

#### X-ray Crystallography

Single-crystal X-ray diffraction analyses were performed using a Rigaku FRE+ equipped with either VHF (**1, 9, 13, 37**) or HF Varimax confocal mirrors (**17, 48, 49**) and an AFC12 goniometer and HG Saturn 724+ detector equipped with an Oxford Cryosystems low-temperature apparatus operating at *T* = 100(2) K. CrystalClear-SM Expert 3.1 b27 (CrystalClear, Rigaku Corporation, The Woodlands, Texas, U.S.A., (2008-2014) (**1, 9, 13, 37**) or CrysAlisPro (CrysAlisPro Software System, Rigaku Oxford Diffraction, Yarnton, Oxford, UK (2016)) (**17, 48, 49**) was used to record images. CrysAlisPro was used to process all data and apply empirical absorption corrections and unit cell parameters were refined against all data. The structures were solved by intrinsic phasing using SHELXT (Sheldrick 2015) (**1, 9, 13, 17, 37, 48**) or charge flipping using SUPERFLIP (Palatinus and Chapuis 2007) (**49**) and refined on *Fo^2^* by full-matrix least-squares refinements using SHELXL-2014 (Sheldrick 2015) as implemented within OLEX2 (Dolomanov, Bourhis et al. 2009). All nonhydrogen atoms were refined with anisotropic displacement parameters and hydrogen atoms were added at calculated positions except those attached to heteroatoms which were located from the difference map. All hydrogen atoms were refined using a riding model with isotropic displacement parameters based on the equivalent isotropic displacement parameter (*Ueq*) of the parent atom. Figures were produced using OLEX2. The CIF files for the crystal structures of **1, 9, 13, 17, 37, 48** and **49** have been deposited with the CCDC and have been given the deposition numbers 1534017-1534023 respectively.

### Compound preparation

**Method A** - Quinoline (200 mg, 0.86 mmol) and aniline (0.95 mmol) were suspended in ethanol (10 mL) and refluxed at 90 °C for 18 hours. The product was collected by filtration and washed with water (2 × 10 mL) and ether (3 × 10 mL) to yield the product as a free following solid.

**Method B** - Quinoline/Quinazoline (200 mg, 0.86 mmol) and aniline (0.95 mmol) were suspended in ethanol (10 mL) and refluxed at 90 °C for 18 hours. The crude mixture was purified by flash chromatography ethyl acetate:hexane followed by 1-5% methanol:ethyl acetate and solvent removed under reduced pressure to yield the product as a free following solid.

**Method C** - Quinazoline (200 mg, 0.864 mmol) and aniline (0.95 mmol) were suspended in butanol (10 mL) and refluxed at 120°C for 18 hours. The crude mixture was purified by flash chromatography 20-100% ethyl acetate:hexane followed by 1-5% methanol:ethyl acetate and solvent removed under reduced pressure to yield the product as a free following solid.

**Method D** - Quinazoline (200 mg, 0.86 mmol) and aniline (0.95 mmol) palladium acetate (20 %, 40 mg, 0.17 mmol), tri(*o*-tolyl)phosphine (20 %, 55 mg, 0.18 mmol) and caesium carbonate (220 %, 381 mg, 2.0 mmol) were all suspended in acetonitrile 10 mL and de-gased for 5 minutes. The mixture was refluxed at 85 °C for 18 hours. The crude mixture was passed through a plug of celite 545 before been purified by flash chromatography 20-100% ethyl acetate:hexane followed by 1-5% methanol:ethyl acetate and solvent removed under reduced pressure to yield the product as a free following solid.

**6-(trifluoromethyl)-*N*-(3,4,5-trimethoxyphenyl)quinolin-4-amine (1)** prepared by method B to afford a mustard solid (219 mg, 67%) ^1^H NMR (400 MHz, DMSO-*d*_6_) δ 11.23 (s, 1H), 9.30 (s, 1H), 8.57 (d, *J* = 7.0 Hz, 1H), 8.29 (s, 2H), 6.96 (d, *J* = 7.0 Hz, 1H), 6.83 (s, 2H), 3.77 (d, *J* = 29.8 Hz, 9H). ^13^C NMR (101 MHz, DMSO-*d*_6_) δ 155.30, 153.69 (s, 2C), 144.56, 140.70, 136.65, 132.59, 129.06, 126.62 (d, *J* = 32.9 Hz, CF_3_), 125.15, 122.46 (d, *J* = 5.1 Hz, 2C), 116.62, 103.07 (s, 2C), 101.54, 60.19, 56.16 (s, 2C); δ HRMS-ESI (m/z): [M+H]^+^ calcd for C1_9_H_18_F_3_N_2_O_3_ - 379.1270; found 379.1261; LC - T_r_ = 3.86 min, purity >95%.

***N*-(3,5-dimethoxyphenyl)-6-(trifluoromethyl)quinolin-4-amine (4)** prepared by method B to afford a tan solid (47 mg, 16%) ^1^H NMR (400 MHz, DMSO-*d*_6_) δ 9.72 (s, 1H), 8.99 (s, 1H), 8.58 (d, *J* = 5.6 Hz, 1H), 8.09 (d, *J* = 8.8 Hz, 1H), 7.98 (dd, *J* = 8.9, 1.7 Hz, 1H), 7.10 (d, *J* = 5.6 Hz, 1H), 6.58 (d, *J* = 2.2 Hz, 2H), 6.37 (t, *J* = 2.2 Hz, 1H), 3.76 (s, 6H); ^13^C NMR (100MHz, DMSO-d_6_) 161.10, 153.68, 149.11, 141.46, 139.45, 135.62, 133.46, 128.32, 125.29, 124.81, 124.47, 121.37, 118.63, 103.06 (s, 2C), 100.96, 55.31 (s, 2C); HRMS-ESI (m/z): [M+H]^+^ calcd for C_18_H_16_F_3_N_2_O_2_ - 349.1164; found 349.1149; LC - T_r_ = 4.22 min, purity >95%.

***N*-(3,4-dimethoxyphenyl)-6-(trifluoromethyl)quinolin-4-amine (5)** prepared by method B to afford a yellow solid (165 mg, 55%) ^1^H NMR (400 MHz, DMSO-*d*_6_) δ 11.25 (s, 1H), 9.32 (s, 1H), 8.54 (d, *J* = 6.9 Hz, 1H), 8.28 (s, 2H), 7.14 (d, *J* = 8.5 Hz, 1H), 7.09 (s, 1H), 7.02 (d, *J* = 8.3 Hz, 1H), 6.81 (d, *J* = 6.9 Hz, 1H), 3.80 (d, *J* = 18.3 Hz, 6H); ^13^C NMR (101 MHz, DMSO-*d*_6_) δ 155.58, 149.57, 148.20, 144.26, 140.61, 129.53, 129.06 (d, *J* = 3.4 Hz), 126.56 (q, *J* = 32.7 Hz, CF_3_), 125.16, 122.56 – 122.09 (m, 2C), 117.63, 116.54, 112.35, 109.63, 101.03, 55.75 (s, 2C). HRMS-ESI (m/z): [M+H]^+^ calcd for C_18_H_16_F_3_N_2_O_2_ −349.1164; found 349.1153; T_r_ = 3.85 min, purity >95%.

***N*-(2,4-dimethoxyphenyl)-6-(trifluoromethyl)quinolin-4-amine (6)** prepared by method B to afford a mustard solid (202 mg, 67%) ^1^H NMR (400 MHz, DMSO-*d*_6_) δ 9.30 (s, 1H), 8.48 (d, *J* = 6.6 Hz, 1H), 8.25 (d, *J* = 8.8 Hz, 1H), 8.17 (dd, *J* = 8.9, 1.4 Hz, 1H), 7.29 (d, *J* = 8.6 Hz, 1H), 6.80 (d, *J* = 2.5 Hz, 1H), 6.67 (dd, *J* = 8.6, 2.6 Hz, 1H), 6.31 (d, *J* = 6.6 Hz, 1H), 3.83 (s, 3H), 3.77 (s, 3H); ^13^C NMR (101 MHz, DMSO-*d*_6_) δ 160.19, 155.39, 154.95, 145.67, 142.49, 128.75, 128.00 (d, *J* = 2.9 Hz), 126.09 (q, *J* = 32.7 Hz, CF_3_), 124.01, 122.64, 122.30 (d, *J* = 4.2 Hz), 118.01, 116.61, 105.61, 101.42, 99.82, 55.83, 55.61; HRMS-ESI (m/z): [M+H]^+^ calcd for C_18_H_16_F_3_N_2_O_2_-349.1164; found 349.1152; T_r_ = 4.14 min, purity >95%.

***N*-(2,5-dimethoxyphenyl)-6-(trifluoromethyl)quinolin-4-amine (7)** prepared by method B to afford a mustard solid (204 mg, 68%) ^1^H NMR (400 MHz, DMSO-*d*_6_) δ 8.34 (s, 1H), 7.72 (d, *J* = 6.7 Hz, 1H), 7.42 – 7.36 (m, 2H), 6.44 (d, *J* = 8.7 Hz, 1H), 6.31 – 6.25 (m, 2H), 5.80 (d, *J* = 6.7 Hz, 1H), 3.04 (d, *J* = 4.1 Hz, 6H); ^13^C NMR (101 MHz, DMSO-*d*_6_) δ 155.42 (d, *J* = 96.8 Hz), 149.46, 146.57, 143.44, 129.08 (d, *J* = 3.1 Hz), 128.26 (q, *J* = 33.0 Hz, CF_3_), 126.65, 126.24, 124.73, 122.55 (d, *J* = 4.3 Hz), 117.91, 114.91, 114.49, 114.22, 102.96, 56.49, 56.15; HRMS-ESI (m/z): [M+H]^+^ calcd for C_18_H_16_F_3_N_2_O_2_ – 349.1164; found 349.1152; T_r_ = 4.05 min, purity >95%.

***N*-(4-methoxyphenyl)-6-(trifluoromethyl)quinolin-4-amine (8)** prepared by method B to afford a green solid (190 mg, 68%); ^1^H NMR (400 MHz, DMSO-*d*_6_) δ 11.27 (s, 1H), 9.33 (d, *J* = 1.7 Hz, 1H), 8.54 (d, *J* = 7.0 Hz, 1H), 8.28 (d, *J* = 1.1 Hz, 2H), 7.41 (d, *J* = 8.9 Hz, 2H), 7.14 (d, *J* = 8.9 Hz, 2H), 6.72 (d, *J* = 7.0 Hz, 1H), 3.83 (s, 3H); ^13^C NMR (101 MHz, DMSO-*d*_6_) δ 158.50, 155.56, 144.28, 140.64, 129.39, 129.06 (d, *J* = 3.4 Hz), 127.00 (s, 2C), 126.54 (q, *J* = 33.0 Hz, CF_3_), 125.15, 122.44, 122.32, 116.57, 115.19 (s, 2C), 100.72, 55.47; HRMS-ESI (m/z): [M+H]^+^ calcd for C_17_H_14_F_3_N_2_O – 319.1058; found 319.1049; T_r_ = 4.04 min, purity >95%.

***N*-(3-methoxyphenyl)-6-(trifluoromethyl)quinolin-4-amine (9)** prepared by method B to afford a light yellow solid (258 mg, 94%) ^1^H NMR (400 MHz, DMSO-*d*_6_) δ 11.24 (s, 1H), 9.34 (s, 1H), 8.58 (d, *J* = 6.8 Hz, 1H), 8.28 (q, *J* = 8.9 Hz, 2H), 7.48 (t, *J* = 8.3 Hz, 1H), 7.07 (s, 2H), 6.96 (dd, *J* = 26.3, 7.3 Hz, 2H), 3.81 (s, 3H); ^13^C NMR (101 MHz, DMSO-*d*_6_) δ 160.55, 154.81, 145.35, 141.64, 138.60, 130.97, 128.95 (d, *J* = 3.4 Hz), 126.70 (q, *J* = 32.8 Hz, CF_3_), 125.41, 123.23, 122.71 (d, *J* = 3.9 Hz), 117.19, 117.16, 113.16, 110.94, 101.63, 55.62; HRMS-ESI (m/z): [M+H]^+^ calcd for C_17_H_14_F_3_N_2_O – 319.1058; found 319.1048; T_r_ = 4.08 min, purity >95%.

***N*-(2-methoxyphenyl)-6-(trifluoromethyl)quinolin-4-amine (10)** prepared by method B to afford a dark green solid (104 mg, 38%) ^1^H NMR (400 MHz, DMSO-*d*_6_) δ 9.35 (s, 1H), 8.52 (d, *J* = 6.7 Hz, 1H), 8.29 (d, *J* = 8.9 Hz, 1H), 8.22 (dd, *J* = 8.9, 1.7 Hz, 1H), 7.46 (ddd, *J* = 8.3, 7.5, 1.7 Hz, 1H), 7.41 (dd, *J* = 7.8, 1.6 Hz, 1H), 7.28 (dd, *J* = 8.4, 1.1 Hz, 1H), 7.12 (td, *J* = 7.6, 1.2 Hz, 1H), 6.36 (d, *J* = 6.7 Hz, 1H), 3.79 (s, 3H); ^13^C NMR (101 MHz, DMSO-*d*_6_) δ 154.64, 154.21, 142.03, 129.39, 128.25 (d, *J* = 2.9 Hz), 127.85 126.25 (d, *J* = 32.7 Hz, CF_3_), 125.29, 125.10, 123.64, 122.57, 122.42 (d, *J* = 4.2 Hz), 121.20, 116.62, 112.89, 101.68, 55.69; HRMS-ESI (m/z): [M+H]^+^ calcd for C_17_H_14_F_3_N_2_O – 319.1058; found 319.1049; T_r_ = 4.00 min, purity >95%.

***N*-(benzo[*d*][1,3]dioxol-5-yl)-6-(trifluoromethyl)quinolin-4-amine (11)** prepared by method B to afford a dark green solid (126 mg, 44%) ^1^H NMR (400 MHz, DMSO-*d*_6_) δ 11.24 (s, 1H), 9.32 (s, 1H), 8.56 (d, *J* = 6.9 Hz, 1H), 8.28 (s, 2H), 7.14 – 7.05 (m, 2H), 6.95 (dd, *J* = 8.2, 2.0 Hz, 1H), 6.78 (d, *J* = 6.9 Hz, 1H), 6.14 (s, 2H); ^13^C NMR (101 MHz, DMSO-*d*_6_) δ 155.60, 148.26, 146.65, 144.43, 140.74, 130.63, 129.00 (d, *J* = 2.7 Hz), 126.54 (q, *J* = 32.8 Hz, CF_3_), 125.15, 122.46 (d, *J* = 3.2 Hz, 2C), 119.15, 116.58, 108.98, 106.83, 101.93, 101.07; HRMS-ESI (m/z): [M+H]^+^ calcd for C_17_H_12_F_3_N_2_O_2_ - 333.0851; found 333.0840; T_r_ = 3.90 min, purity >95%.

***N*-(2,3-dihydrobenzo[*b*][1,4]dioxin-6-yl)-6-(trifluoromethyl)quinolin-4-amine (12)** prepared by method B to afford a mustard solid (185 mg, 62%) ^1^H NMR (400 MHz, DMSO-*d*_6_) δ 11.33 (s, 1H), 9.36 (s, 1H), 8.55 (d, *J* = 7.0 Hz, 1H), 8.33 – 8.24 (m, 2H), 7.05 (d, *J* = 8.5 Hz, 1H), 7.01 (d, *J* = 2.4 Hz, 1H), 6.94 (dd, *J* = 8.5, 2.5 Hz, 1H), 6.78 (d, *J* = 7.0 Hz, 1H), 4.31 (s, 4H); ^13^C NMR (101 MHz, DMSO-*d*_6_) δ 155.55, 144.14, 144.01, 142.89, 129.90, 129.10 (d, *J* = 2.8 Hz), 127.86, 126.56 (q, *J* = 32.8 Hz, CF_3_), 122.62 (d, *J* = 4.5 Hz), 122.43, 122.17, 118.50, 118.07, 116.56, 114.42, 100.93, 64.13 (s, 2C). HRMS-ESI (m/z): [M+H]^+^ calcd for C_18_H_14_F_3_N_2_O_2_ – 347.1007; found 347.0997; T_r_ = 4.05 min, purity >95%.

**6-(trifluoromethyl)-*N*-(3,4,5-trimethoxyphenyl)quinazolin-4-amine (13)** prepared by method C to afford a yellow solid (137 mg, 42%) ^1^H NMR (400 MHz, DMSO-*d*_6_) δ 11.41 (s, 1H), 9.26 (s, 1H), 8.94 (s, 1H), 8.33 (dd, *J* = 8.8, 1.6 Hz, 1H), 8.07 (d, *J* = 8.7 Hz, 1H), 7.14 (s, 2H), 3.81 (s, 6H). 3.71 (s, 3H). ^13^C NMR (101 MHz, DMSO-*d*_6_) δ 153.70, 152.74 (s, 2C), 135.92, 134.53, 132.64, 130.99, 130.90, 129.40, 127.80, 126.52, 122.85, 122.29, 113.76, 102.33 (s, 2C), 60.20, 56.07 (s, 2C). HRMS-ESI (m/z): [M+H]^+^ calcd for C_18_H_14_F_3_N_2_O_2_ – 380.1222; found 380.1210; T_r_ = 4.38 min, purity >95%.

***N*-(3,5-dimethoxyphenyl)-6-(trifluoromethyl)quinazolin-4-amine (14)** prepared by method D to afford a light yellow solid (114 mg, 38%) ^1^H NMR (400 MHz, DMSO-*d*_6_) δ 11.51 (s, 1H), 9.32 (s, 1H), 8.98 (s, 1H), 8.34 (d, *J* = 10.3 Hz, 1H), 8.11 (d, *J* = 8.7 Hz, 1H), 7.01 (s, 2H), 6.50 (t, *J* = 2.2 Hz, 1H), 3.79 (s, 6H); ^13^C NMR (101 MHz, DMSO-*d*_6_) δ 160.87 (s, 2C), 160.02, 153.86, 138.80-131.54 (m, 1C, CF_3_), 128.37, 128.03, 125.41, 123.54-123.43 (m, 1C), 122.70, 120.52, 114.22, 103.23 (s, 2C), 98.60, 55.88 (s, 2C); HRMS-ESI (m/z): [M+H]^+^ calcd for C1_8_H_15_F_3_N_3_O_2_ - 350.1116; found 350.1110; T_r_ = 4.84 min, purity >95%.

***N*-(4-methoxyphenyl)-6-(trifluoromethyl)quinazolin-4-amine (15)** prepared by method D to afford a light yellow solid (121 mg, 44%) ^1^H NMR (400 MHz, DMSO-*d*_6_) δ 11.86 (s, 1H), 9.37 (s, 1H), 8.93 (s, 1H), 8.36 (dd, *J* = 8.8, 1.6 Hz, 1H), 8.12 (d, *J* = 8.7 Hz, 1H), 7.77 – 7.51 (m, 2H), 7.19 – 6.95 (m, 2H), 3.81 (s, 3H); ^13^C NMR (101 MHz, DMSO-*d*_6_) δ 159.44, 157.89, 155.76, 152.84, 131.38, 129.15 (d, *J* = 3.7 Hz), 128.05, 126.08 (**s**, 2C), 123.29-123.13 (m, 1C), 122.22, 121.94, 114.02, 113.58 (**s**, 2C), 55.40; HRMS-ESI (m/z): [M+H]^+^ calcd for C_16_H_13_F_3_N_3_O - 320.1011; found 320.1004; T_r_ = 4.08 min, purity >95%.

**6-(trifluoromethyl)-*N*-(3,4,5-trifluorophenyl)quinolin-4-amine (16)** prepared by method B to afford a tan solid (142 mg, 48%) ^1^H NMR (400 MHz, DMSO-*d*_6_) δ 9.32 (s, 1H), 8.65 (d, *J* = 5.2 Hz, 1H), 8.24 (dd, *J* = 39.6, 8.5 Hz, 2H), 7.70 – 7.37 (m, 2H), 7.09 (d, *J* = 5.6 Hz, 1H). ^13^C NMR (101 MHz, DMSO-*d*_6_) δ 153.02, 151.85 (dd, *J* = 10.3, 4.7 Hz), 149.39 (dd, *J* = 9.8, 5.5 Hz), 146.96, 143.12, 138.18 (t, *J* = 14.7 Hz), 135.71 (t, *J* = 15.5 Hz), 134.70 (td, *J* = 10.8, 3.5 Hz), 128.09 – 127.92 (m, 1C), 126.38 (q, *J* = 32.7 Hz, CF_3_), 125.24, 124.56, 122.53, 117.63, 109.45 (d, *J* = 23.1 Hz), 102.51; HRMS-ESI (m/z): [M+H]^+^ calcd for C_16_H_9_N_2_F_6_ - 343.0670; found 343.0653; Full Scan at T_r_ = 4.24 min, purity >95%.

***N*-(3,5-difluorophenyl)-6-(trifluoromethyl)quinolin-4-amine (17)** prepared by method B to afford an off white solid (169 mg, 64%) ^1^H NMR (400 MHz, DMSO-*d*_6_) δ 11.64 (s, 1H), 9.42 (s, 1H), 8.69 (d, *J* = 7.0 Hz, 1H), 8.37 (d, *J* = 9.0 Hz, 1H), 8.33 (dd, *J* = 9.0, 1.6 Hz, 1H), 7.52 – 7.22 (m, 3H), 7.15 (d, *J* = 7.0 Hz, 1H); ^13^C NMR (101 MHz, DMSO-*d*_6_) δ 164.11 (d, *J* = 15.0 Hz), 161.66 (d, *J* = 15.1 Hz), 155.06, 144.69, 140.21, 139.84 (t, *J* = 13.1 Hz), 130.26 – 128.72 (m, 1C), 127.04 (q, *J* = 32.9 Hz), 125.04, 122.78 (q, *J* = 3.7 Hz), 122.33, 122.17, 117.03, 109.62 – 107.92 (m, 1C), 103.24, 102.98 (t, *J* = 26.0 Hz); HRMS-ESI (m/z): [M+H]^+^ calcd for C16H_10_N_2_F_5_ - 325.0764; 325.07487; T_r_ = 4.07 min, purity >95%.

***N*-(4-fluorophenyl)-6-(trifluoromethyl)quinolin-4-amine (18)** prepared by method B to afford an off white solid (169 mg, 64%) ^1^H NMR (400 MHz, DMSO-*d*_6_) δ 11.57 (s, 1H), 9.48 (s, 1H), 8.57 (d, *J* = 6.9 Hz, 1H), 8.36 (d, *J* = 8.9 Hz, 1H), 8.28 (dd, *J* = 8.9, 1.4 Hz, 1H), 7.68 – 7.48 (m, 2H), 7.48 – 7.31 (m, 2H), 6.78 (d,J = 6.9 Hz, 1H); ^13^C NMR (101 MHz, DMSO-*d*_6_) δ 162.03, 159.60, 155.36, 144.48, 140.80, 133.42, 129.01 (d, *J* = 3.1 Hz), 127.77 (d, *J* = 8.7 Hz), 126.61 (q, *J* = 32.8 Hz), 125.16, 122.88 (d, *J* = 4.2 Hz), 122.39, 116.80. (d, *J* = 8.7 Hz), 116.78, 100.93; HRMS-ESI (m/z): [M+H]^+^ calcd for C1_6_H_11_N_2_F_4_ - 307.0858; found 307.0844; T_r_ = 3.86 min, purity >95%.

***N*-(3-fluorophenyl)-6-(trifluoromethyl)quinolin-4-amine (19)** prepared by method B to afford a light yellow solid (204 mg, 77%) ^1^H NMR (400 MHz, DMSO-*d*_6_) δ 11.38 (s, 1H), 9.37 (s, 1H), 8.62 (d, *J* = 6.6 Hz, 1H), 8.32 (d, *J* = 8.8 Hz, 1H), 8.24 (d, *J* = 8.5 Hz, 1H), 7.59 (q, *J* = 7.9 Hz, 1H), 7.38 (dd, *J* = 15.9, 9.0 Hz, 2H), 7.23 (td, *J* = 8.5, 2.0 Hz, 1H), 7.01 (d, *J* = 6.6 Hz, 1H); ^13^C NMR (101 MHz, DMSO-*d*_6_) δ 163.76, 161.32, 153.87, 145.92, 142.11, 139.41 (d, *J* = 10.3 Hz), 131.50 (d, *J* = 9.4 Hz), 128.46 (d, *J* = 3.0 Hz), 126.45 (q, *J* = 32.7 Hz, CF_3_), 123.62, 122.85 – 121.96 (m, 1C), 120.69 (d, *J* = 2.8 Hz), 117.27, 113.61 (d, *J* = 20.9 Hz), 111.81 (d, *J* = 23.8 Hz), 101.78; HRMS-ESI (m/z): [M+H]^+^ calcd for C1_6_H_11_N_2_F_4_ 307.0858; found 307.0844; T_r_ = 3.95 min, purity >95%.

***N*-(2-fluorophenyl)-6-(trifluoromethyl)quinolin-4-amine (20)** prepared by method B to afford a light yellow solid (135 mg, 51%) ^1^H NMR (400 MHz, DMSO-*d*_6_) δ 9.37 (s, 1H), 8.61 (d, *J* = 6.4 Hz, 1H), 8.35 – 8.16 (m, 2H), 7.58 (t, *J* = 7.8 Hz, 1H), 7.54 – 7.44 (m, 2H), 7.39 (dt, *J* = 8.2, 4.7 Hz, 1H), 6.54 (dd, *J* = 6.3, 2.3 Hz, 1H); ^13^C NMR (101 MHz, DMSO-*d*_6_) δ 157.91, 155.43, 153.97, 146.53, 142.69, 129.44 (d, *J* = 7.8 Hz), 128.50, 128.13 – 128.08 (m, 1C), 126.37 (q, *J* = 32.8 Hz), 125.70 (d, *J* = 3.6 Hz), 125.26, 125.02 (d, *J* = 12.1 Hz), 124.32, 122.56 – 122.37 (m, 1C), 117.17 – 116.64 (m), 101.72; HRMS-ESI (m/z): [M+H]^+^ calcd for C1_6_H_11_N_2_F_4_ −307.0858;found 307.0844; T_r_ = 3.81 min, purity >95%.

***N*-(4-chloro-3-fluorophenyl)-6-(trifluoromethyl)quinolin-4-amine (21)** prepared by method B to afford a bright yellow solid (200 mg, 68%) ^1^H NMR (400 MHz, DMSO-*d*_6_) δ ^1^H NMR (400 MHz, DMSO-*d*_6_) δ 9.26 (s, 1H), 8.63 (d, *J* = 5.8 Hz, 1H), 8.19 (dd, *J* = 44.3, 8.6 Hz, 2H), 7.69 (t, *J* = 8.4 Hz, 1H), 7.64 – 7.53 (m, 1H), 7.37 (d, *J* = 8.2 Hz, 1H), 7.10 (d, *J* = 5.9 Hz, 1H); ^13^C NMR (101 MHz, DMSO-*d*_6_) δ 158.64, 156.19, 151.67, 148.23 (d, *J* = 1.6 Hz), 144.77 (d, *J* = 1.6 Hz), 139.47 (d, *J* = 9.3 Hz), 131.29, 127.28, 125.12 (q, *J* = 33.3 Hz, CF_3_), 125.89, 122.27 (q, *J* = 3.7 Hz), 120.71 (d, *J* = 2.9 Hz), 118.03, 115.67 (d, *J* = 17.6 Hz), 112.04 (d, *J* = 23.4 Hz), 102.67; HRMS-ESI (m/z): [M+H]^+^ calcd for C16H_10_N_2_F4CI – 341.0469; found 341.0454; T_r_ = 4.32 min, purity >95%.

***N*-(3-chloro-5-fluorophenyl)-6-(trifluoromethyl)quinolin-4-amine (22)** prepared by method B to afford a tan solid (185 mg, 63%) ^1^H NMR (400 MHz, DMSO-*d*_6_) δ 9.19 (s, 1H), 8.66 (s, 1H), 8.18 (dd, *J* = 34.8, 7.1 Hz, 2H), 7.41 (s, 1H), 7.33 (dd, *J* = 22.7, 8.5 Hz, 2H), 7.15 (s, 1H); ^13^C NMR (101 MHz, DMSO-*d*_6_) δ 163.76, 161.30, 151.69, 147.99, 144.19, 141.51 (d, *J* = 11.8 Hz), 134.66 (d, *J* = 12.9 Hz), 127.45, 126.14 (q, *J* = 32.4 Hz), 125.52, 122.11 (d, *J* = 3.9 Hz), 119.40 - 119.35 (m, 1C), 118.00, 112.85 (d, *J* = 25.2 Hz), 109.26 (d, *J* = 24.1 Hz), 103.07; HRMS-ESI (m/z): [M+H]^+^ calcd for C_16_H_10_N_2_F_4_CI – 341.0469; found 341.0458; T_r_ = 4.40 min, purity >95%.

***N*-(3,4-dichlorophenyl)-6-(trifluoromethyl)quinolin-4-amine (23)** prepared by method B to afford a tan solid (207 mg, 67%) ^1^H NMR (400 MHz, DMSO-*d*_6_) δ ^1^H NMR (400 MHz, DMSO-*d_6_*) δ 9.20 (s, 1H), 8.71 – 8.51 (m, 1H), 8.15 (dd, *J* = 42.3, 8.2 Hz, 2H), 7.77 – 7.66 (m, 2H), 7.48 (d, *J* = 7.4 Hz, 1H), 7.06 (d, *J* = 4.4 Hz, 1H). ^13^C NMR (101 MHz, DMSO-*d*_6_) δ 151.34, 148.59, 145.15, 139.16, 131.73, 131.33, 127.09 – 126.75 (m, 1C), 126.95, 126.25, 125.84 (q, *J* = 32.3 Hz), 125.08, 123.54, 122.65, 122.08 (d, *J* = 3.9 Hz), 118.07, 102.59; HRMS-ESI (m/z): [M+H]^+^ calcd for C_16_H_10_N_2_F_3_CI_2_ - 357.0173; found 357.0163; T_r_ = 4.71 min, purity >95%.

***N*-(4-chlorophenyl)-6-(trifluoromethyl)quinolin-4-amine (24)** prepared by method B to afford a light yellow solid (181 mg, 65%) ^1^H NMR (400 MHz, DMSO-*d*_6_) δ 9.28 (s, 1H), 8.58 (d, *J* = 6.3 Hz, 1H), 8.24 (d, *J* = 8.8 Hz, 1H), 8.14 (dd, *J* = 8.9, 1.6 Hz, 1H), 7.59 – 7.48 (m, 4H), 6.94 (d, *J* = 6.3 Hz, 1H); ^13^C NMR (101 MHz, DMSO-*d*_6_) δ 152.36, 147.88, 144.55, 137.42, 129.94, 129.64 (s, 2C), 128.15 (d, *J* = 2.5 Hz), 127.48 – 127.03 (m, 1C), 126.44 – 125.32 (m, 3C), 122.72, 122.28 (d, *J* = 4.4 Hz), 117.77, 101.80; HRMS-ESI (m/z): [M+H]^+^ calcd for C1_6_H_11_N_2_F_3_CI-323.0563;found 323.0553; T_r_ = 4.30 min, purity >95%.

***N*-(3-chlorophenyl)-6-(trifluoromethyl)quinolin-4-amine (25)** prepared by method B to afford a light yellow solid (201 mg, 72%) ^1^H NMR (400 MHz, DMSO-*d*_6_) δ 11.40 (s, 1H), 9.35 (s, 1H), 8.62 (d, *J* = 6.6 Hz, 1H), 8.41 – 8.19 (m, 2H), 7.61 (s, 1H), 7.57 (d, *J* = 7.9 Hz, 1H), 7.47 (dd, *J* = 14.7, 7.8 Hz, 2H), 6.98 (d, *J* = 6.6 Hz, 1H); ^13^C NMR (101 MHz, DMSO-*d*_6_) δ 154.03, 145.81, 141.91, 139.14, 133.91, 131.46, 128.57 (d, *J* = 3.1 Hz), 126.80, 126.51 (q, *J* = 32.3Hz, CF_3_), 125.19, 124.60, 123.49, 122.77 – 122.30 (m), 122.48, 117.24, 101.71; HRMS-ESI (m/z): [M+H]^+^ calcd for C_16_H_11_N_2_F_3_CI – 323.0563; found 323.0553; T_r_ = 4.25 min, purity >95%.

***N*-(2-chlorophenyl)-6-(trifluoromethyl)quinolin-4-amine (26)** prepared by method B to afford a light yellow solid (187 mg, 67%) ^1^H NMR (400 MHz, DMSO-*d*_6_) δ 9.42 (s, 1H), 8.61 (s, 1H), 8.26 (dd, *J* = 36.5, 7.3 Hz, 2H), 7.84 – 7.34 (m, 4H), 6.96 (s, 1H); ^13^C NMR (101 MHz, DMSO-*d*_6_) δ 154.45, 146.15, 142.30, 139.60, 134.30, 131.85, 129.06 – 128.85 (m, 1C), 127.15, 126.93 (q, *J* = 32.9 Hz, CF_3_), 125.60, 125.01, 123.79, 123.37 – 123.05 (m, 1C), 122.89, 117.68, 102.15; HRMS-ESI (m/z): [M+H]^+^ calcd for C_16_H_11_N_2_F_3_CI – 323.0563; found 323.0553; T_r_ = 4.04 min, purity >95%.

***N*-(4-bromophenyl)-6-(trifluoromethyl)quinolin-4-amine (27)** prepared by method B to afford a yellow solid (225 mg, 71%) ^1^H NMR (400 MHz, DMSO-*d*_6_) δ 9.27 (s, 1H), 8.58 (d, *J* =4.7 Hz, 1H), 8.19 (dd, *J* = 36.2, 8.4 Hz, 2H), 7.56 (dd, *J* = 100.3, 7.7 Hz, 4H), 6.95 (d, *J* = 5.0 Hz, 1H); ^13^C NMR (101 MHz, DMSO-*d*_6_) δ 152.48, 147.48, 144.03, 137.63, 132.55, 127.57 – 127.39 (m, 1C), 126.25, 125.95 (d, *J* = 32.6 Hz, CF_3_), 125.32, 125.24, 122.61, 122.23 (d, *J* = 4.1 Hz), 118.28, 117.64, 101.75; HRMS-ESI (m/z): [M+H]^+^ calcd for C_16_H_11_N_2_F_3_Br – 367.0058; found 367.0046; T_r_ = 4.37 min, purity >95%.

***N*-(3-bromophenyl)-6-(trifluoromethyl)quinolin-4-amine (28)** prepared by method B to afford a yellow solid (251 mg, 79%) ^1^H NMR (500 MHz, DMSO-*d*_6_) δ 9.28 (s, 1H), 8.61 (d, *J* = 6.1 Hz, 1H), 8.27 – 8.17 (m, 2H), 7.70 (s, 1H), 7.51 (q, *J* = 11.9, 9.9 Hz, 3H), 6.99 (d, *J* = 6.1 Hz, 1H); ^13^C NMR (126 MHz, DMSO-*d*_6_) δ 153.07, 146.95, 143.22, 139.75, 131.65, 129.17, 128.18 – 127.70 (m, 1C), 126.97, 126.27 (q, *J* = 32.6 Hz, CF_3_), 125.03, 124.62, 123.28, 122.86, 122.36-122.21 (m, 1C), 117.54, 101.94; HRMS-ESI (m/z): [M+H]^+^ calcd for C_16_H_11_N_2_F_3_Br - 367.0058; found 367.0046; T_r_ = 4.39 min, purity >95%.

***N*-(2-bromophenyl)-6-(trifluoromethyl)quinolin-4-amine (29)** prepared by method B to afford a yellow solid (219 mg, 69%) ^1^H NMR (400 MHz, DMSO-*d*_6_) δ 9.09 (s, 1H), 8.46 (d, *J* = 5.8 Hz, 1H), 8.14 – 7.98 (m, 2H), 7.83 (dd, *J* = 8.1, 1.2 Hz, 1H), 7.62 – 7.45 (m, 2H), 7.34 (td, *J* = 7.6, 1.8 Hz, 1H), 6.18 (d, *J* = 5.8 Hz, 1H); ^13^C NMR (101 MHz, DMSO-*d*_6_) δ 151.78, 150.26, 147.57, 138.54, 134.16, 129.78, 129.26, 128.35, 126.39 – 126.15 (m, 1C), 126.10, 125.56 (q, *J* = 32.2 Hz), 123.40, 122.16 – 121.72 (m, 2C), 118.33, 102.47; HRMS-ESI (m/z): [M+H]^+^ calcd for C_16_H_11_N_2_F_3_Br – 367.0058; found 367.0046; T_r_ = 4.10 min, purity >95%.

***N*-(3-iodophenyl)-6-(trifluoromethyl)quinolin-4-amine (30)** prepared by method B to afford a yellow solid (240 mg, 67%) ^1^H NMR (400 MHz, DMSO-*d*_6_) δ 9.22 (s, 1H), 8.58 (d, *J* = 6.2 Hz, 1H), 8.23 – 8.08 (m, 2H), 7.81 (s, 1H), 7.57 (dd, *J* = 60.3, 8.3 Hz, 2H), 7.28 (t, *J* = 7.9 Hz, 1H), 6.95 (d, *J* = 6.2 Hz, 1H); ^13^C NMR (101 MHz, DMSO-*d*_6_) δ 152.23, 147.81, 144.32, 139.88, 134.55, 132.34, 131.50, 127.38 (d, *J* = 3.0 Hz), 125.96 (q, *J* = 32.5 Hz, CF_3_), 125.51, 123.25, 122.67, 122.26, 122.20 (q, *J* = 4.1 Hz), 122.14, 117.74, 101.97, 95.30; HRMS-ESI (m/z): [M+H]^+^ calcd for C_16_H_11_N_2_F_3_l – 414.9919; found 414.9902; T_r_ = 4.55 min, purity >95%.

**4-((6-(trifluoromethyl)quinolin-4-yl)amino)benzonitrile (31)** prepared by method B to afford a yellow solid (251 mg, 79%) ^1^H NMR (400 MHz, DMSO-*d*_6_) δ 11.76 (s, 1H), 9.45 (s, 1H), 8.71 (d, *J* = 6.9 Hz, 1H), 8.40 – 8.30 (m, 2H), 8.07 – 7.72 (m, 4H), 7.16 (d, *J* = 6.9 Hz, 1H); ^13^C NMR (101 MHz, DMSO-*d*_6_) δ 154.65, 144.78, 141.75, 140.40, 134.03 (s, 2C), 129.42 (d, *J* = 3.1 Hz), 127.02 (q, *J* = 32.9 Hz), 125.17 (s, 2C), 122.94 (q, *J* = 4.0 Hz), 122.34, 122.24, 118.48, 117.40, 109.11, 102.18. HRMS-ESI (m/z): [M+H]^+^ calcd for C_17_H_11_N_3_F_3_ - 314.0905; found 314.0892; T_r_ = 3.59 min, purity >95%.

**3-((6-(trifluoromethyl)quinolin-4-yl)amino)benzonitrile (32)** prepared by method B to afford an off-white solid (251 mg, 79%) ^1^H NMR (400 MHz, DMSO-*d*_6_) δ 9.28 (s, 1H), 8.63 (d, *J* = 6.2 Hz, 1H), 8.30 – 8.15 (m, 2H), 7.95 (s, 1H), 7.83 (d, *J* = 7.7 Hz, 1H), 7.80 – 7.66 (m, 2H), 7.06 (d, *J* = 6.1 Hz, 1H); ^13^C NMR (101 MHz, dmso) δ 152.16, 147.84, 144.22, 139.57, 131.06, 129.37, 128.71, 127.66 – 127.33 (m, 1C) 127.02, 126.11 (q, *J* = 32.6 Hz), 125.47, 125.35, 122.63, 122.27 (d, *J* = 4.1 Hz), 117.89, 112.51, 102.31; HRMS-ESI (m/z): [M+H]^+^ calcd for C_17_H_11_N_3_F_3_ - 314.0905; found 314.0892; T_r_ = 3.57 min, purity >95%.

**2-((6-(trifluoromethyl)quinolin-4-yl)amino)benzonitrile (33)** prepared by method B to afford a yellow solid (184 mg, 68%) ^1^H NMR (400 MHz, DMSO-*d*_6_) δ ^1^H NMR (400 MHz, DMSO-*d_6_*) δ 8.84 (s, 1H), 8.30 (s, 1H), 7.95 (s, 2H), 7.88 (d, *J* = 7.8 Hz, 1H), 7.74 (t, *J* = 8.3 Hz, 1H), 7.43 (d, *J* = 8.0 Hz, 1H), 7.35 (t, *J* = 7.9 Hz, 1H), 6.37 (s, 1H); ^13^C NMR (101 MHz, DMSO-*d*_6_) δ ^13^C NMR (101 MHz, DMSO-*d*_6_) δ 154.19, 150.83, 147.02, 140.27, 135.06, 134.43, 131.28, 129.27, 128.96, 126.26, 123.56, 121.72, 117.87, 112.45, 107.95, 103.65, 102.91; HRMS-ESI (m/z): [M+H]^+^ calcd for C_17_H_11_N_3_F_3_ - 314.0905; found 314.0893; T_r_ = 3.31 min, purity >95%.

**6-(trifluoromethyl)-*N*-(3-(trifluoromethyl)phenyl)quinolin-4-amine (34)** prepared by method B to afford a yellow solid (120 mg, 39%) ^1^H NMR (400 MHz, DMSO-*d*_6_) δ 9.17 (s, 1H), 8.62 (s, 1H), 8.14 (dd, *J* = 41.7, 7.8 Hz, 2H), 7.78 (s, 2H), 7.71 (s, 1H), 7.59 (d, *J* = 6.2 Hz, 1H), 7.09 – 6.99 (m, 1H); ^13^C NMR (101 MHz, DMSO-*d*_6_) δ 151.08, 149.14, 145.72, 139.88, 130.79, 130.30 (q, *J* = 31.9 Hz, CF3), 126.96, 126.75 (s, 2C), 125.74 (q, *J* = 32.4 Hz), 125.30 (d, *J* = 23.7 Hz), 122.59 (d, *J* = 23.6 Hz), 122.03 – 121.16 (m, 2C), 119.83 (d, *J* = 3.5 Hz), 118.15, 102.29; HRMS-ESI (m/z): [M+H]^+^ calcd for C_17_H_10_N_2_F_6_ - 356.0748; found 357.0814; T_r_ = 4.60 min, purity >95%.

***N*-(3-ethynylphenyl)-6-(trifluoromethyl)quinolin-4-amine (35)** prepared by method B to afford a mustard solid (159 mg, 59%) ^1^H NMR (500 MHz, DMSO-*d*_6_) δ 9.26 (s, 1H), 8.59 (d, *J* = 6.4 Hz, 1H), 8.25 – 8.14 (m, 2H), 7.58 – 7.41 (m, 4H), 6.94 (d, *J* = 6.4 Hz, 1H), 4.31 (s, 1H); ^13^C NMR (126 MHz, DMSO-*d*_6_) δ 152.97, 147.26, 143.66, 138.49, 130.30, 129.60, 127.94 – 127.58 (m, 1C), 127.28, 126.16 (d, *J* = 32.6 Hz), 125.11, 125.03, 124.96, 123.18, 122.94, 122.29(d, *J* = 4.1 Hz), 117.60, 101.80, 82.70, 81.80; HRMS-ESI (m/z): [M+H]^+^ calcd for C_18_H_12_N_2_F_3_ - 313.0953; found 313.0942; T_r_ = 4.17 min, purity >95%.

***N*-(3-((6-(trifluoromethyl)quinolin-4-yl)amino)phenyl)ethan-1-one (36)** prepared by method B to afford a tan solid (185 mg, 65%) ^1^H NMR (400 MHz, DMSO-*d*_6_) δ 9.39 (s, 1H), 8.59 (d, *J* = 6.4 Hz, 1H), 8.33 – 8.18 (m, 2H), 8.03 (s, 1H), 7.94 (d, *J* = 7.5 Hz, 1H), 7.77 (d, *J* = 7.7 Hz, 1H), 7.69 (t, *J* = 7.7 Hz, 1H), 6.95 (d, *J* = 6.5 Hz, 1H), 2.62 (s, 3H); ^13^C NMR (101 MHz, DMSO-*d*_6_) δ 197.37, 153.67, 146.33, 142.71, 138.31, 130.28, 129.10, 128.32 – 127.95 (m, 1C), 126.97 – 125.80 (m, 2C), 125.29, 124.07, 123.98, 122.60, 122.84 – 122.33 (m), 117.42, 101.48, 26.89; HRMS-ESI (m/z): [M+H]^+^ calcd for C_18_H_14_N_2_OF_3_ - 331.1058; found 331.1047; T_r_ = 3.66 min, purity >95%.

***N*-(3-(methylsulfonyl)phenyl)-6-(trifluoromethyl)quinolin-4-amine (37)** prepared by method B to afford a yellow solid (269 mg, 85%) ^1^H NMR (400 MHz, DMSO-*d*_6_) δ 11.63 (s, 1H), 9.40 (s, 1H), 8.68 (d, *J* = 7.0 Hz, 1H), 8.40 – 8.31 (m, 2H), 8.07 (t, *J* = 1.7 Hz, 1H), 7.98 (dt, *J* = 7.4, 1.5 Hz, 1H), 7.94 – 7.84 (m, 2H), 7.06 (d, *J* = 7.0 Hz, 1H), 3.32 (s, 3H); ^13^C NMR (101 MHz, DMSO-*d*_6_) δ 155.12, 144.71, 142.43, 140.33, 138.02, 131.26, 129.94, 129.41 (d, *J* = 3.1 Hz), 126.96 (q, *J* = 32.9 Hz), 125.62, 125.08, 123.31, 122.69 (q, *J* = 4.0 Hz), 122.37, 122.23, 117.05, 101.41, 43.41; HRMS-ESI (m/z): [M+H]^+^ calcd for C_17_H_14_N_2_O_2_SF_3_ - 367.0728; found 367.0716; T_r_ = 3.27 min, purity >95%.

***N*-(4-(methylsulfonyl)phenyl)-6-(trifluoromethyl)quinolin-4-amine (38)** prepared by method A to afford a tan solid (285 mg, 90%)^1^H NMR (400 MHz, DMSO-*d*_6_) δ 11.75 (s, 1H), 9.47 (s, 1H), 8.70 (d, *J* = 7.0 Hz, 1H), 8.35 (q, *J* = 8.2, 7.4 Hz, 2H), 7.96 (dd, *J* = 119.9, 8.7 Hz, 4H), 7.15 (d, *J* = 7.0 Hz, 1H), 3.30 (s, 3H); ^13^C NMR (101 MHz, DMSO-*d*_6_) δ 154.83, 144.74, 141.94, 140.40, 138.77, 129.42 (d, *J* = 3.1 Hz), 128.88 (s, 2C), 127.01 (q, *J* = 32.9 Hz), 125.15 (s, 2C), 122.93 (q, *J* = 4.3, 3.8 Hz), 122.36 122.23, 117.33, 101.99, 43.54; HRMS-ESI (m/z): [M+H]^+^ calcd for C_17_H_14_N_2_O_2_SF_3_ - 367.0728; found 367.0710; T_r_ = 3.29 min, purity >95%.

***N*-(4-(*tert*-butoxy)phenyl)-6-(trifluoromethyl)quinolin-4-amine (39)** prepared by method B to afford a tan solid (233 mg, 75%) ^1^H NMR (400 MHz, DMSO-*d*_6_) δ 10.95 (s, 1H), 9.31 (s, 1H), 8.53 (d, *J* = 6.5 Hz, 1H), 8.25 (d, *J* = 8.8 Hz, 1H), 8.16 (dd, *J* = 8.9, 1.4 Hz, 1H), 7.37 (d, *J* = 8.7 Hz, 2H), 7.12 (d, *J* = 8.7 Hz, 2H), 6.81 (d, *J* = 6.5 Hz, 1H), 1.34 (s, 9H); ^13^C NMR (101 MHz, DMSO-*d*_6_) δ 153.62, 146.58, 143.29, 132.64, 127.86 – 127.59 (m, 1C), 125.97 (q, *J* = 32.6 Hz, CF3), 125.67, 125.35, 124.59, 124.50, 122.64, 122.34 (q, *J* = 4.0 Hz), 117.20, 101.06, 78.39, 28.52 (s, 3C, ^t^Bu); HRMS-ESI (m/z): [M+H]^+^ calcd for C_20_H_20_N_2_OF_3_ - 361.1528; found 361.1515; T_r_ = 4.86 min, purity >95%.

***N*-(4-((methylsulfonyl)methyl)phenyl)-6-(trifluoromethyl)quinolin-4-amine (40)** prepared by method B to afford a tan solid (210 mg, 64%) ^1^H NMR (400 MHz, DMSO-*d*_6_) δ 10.78 (s, 1H), 9.22 (s, 1H), 8.60 (d, *J* = 6.5 Hz, 1H), 8.26 – 8.14 (m, 2H), 7.65 – 7.43 (m, 4H), 6.99 (d, *J* = 6.4 Hz, 1H), 4.57 (s, 2H), 2.96 (s, 3H); ^13^C NMR (101 MHz, DMSO-*d*_6_) δ 152.89, 147.25, 138.15, 132.35 (s, 2C), 127.81 - 127.74 (m, 1C), 127.15, 126.12 (d, *J* = 32.5 Hz), 125.38, 125.07-124.99 (m, 1C), 124.24 (s, 2C), 122.66, 122.09 (d, *J* = 3.7 Hz), 117.55, 101.75, 58.86, 48.59; HRMS-ESI (m/z): [M+H]^+^ calcd for C_18_H_16_N_2_O_2_SF_3_ - 381.0885; found 381.0874; T_r_ = 3.37 min, purity >95%.

***N*-(3,4,5-trimethoxyphenyl)quinolin-4-amine (41)** Prepared by method B to afford a tan solid (288 mg, 76%) ^1^H NMR (400 MHz, DMSO-*d*_6_) δ 10.84 (s, 1H), 8.82 (d, *J* = 8.4 Hz, 1H), 8.47 (d, *J* = 6.8 Hz, 1H), 8.11 (d, *J* = 8.0 Hz, 1H), 7.98 (t, *J* = 8.0 Hz, 1H), 7.76 (t, *J* = 8.1 Hz, 1H), 6.88 (d, *J* = 6.8 Hz, 1H), 6.82 (s, 2H), 3.80 (s, 6H), 3.72 (s, 3H); ^13^C NMR (101 MHz, DMSO-*d*_6_) δ 154.45, 153.59 (s, 2C), 143.31, 139.28, 136.29, 133.31 (s, 2C), 133.23, 126.66, 123.56, 121.07, 117.26, 103.06, 100.38, 60.18, 56.13 (s, 2C); HRMS-ESI (m/z): [M+H]^+^ calcd for C_18_H_19_N_2_O_3_ - 311.1396; found 311.1387; T_r_ = 3.23 min, purity >95%.

**6-fluoro-*N*-(3,4,5-trimethoxyphenyl)quinolin-4-amine (42)** Prepared by method A to afford a light greed solid (288 mg, 76%) ^1^H NMR (400 MHz, DMSO-*d*_6_) δ 10.88 (s, 1H), 8.72 (dd, *J* = 10.5, 2.7 Hz, 1H), 8.50 (d, *J* = 6.9 Hz, 1H), 8.20 (dd, *J* = 9.3, 5.1 Hz, 1H), 7.98 (ddd, *J* = 9.3, 8.0, 2.7 Hz, 1H), 6.92 (d, *J* = 6.9 Hz, 1H), 6.82 (s, 2H), 3.80 (s, 6H), 3.72 (s, 3H); ^13^C NMR (101 MHz, dmso) δ 160.98, 158.53, 154.72, 153.68 (s, 2C), 142.64, 136.59, 135.48, 132.71, 123.491-23.16 (m, 1C), 118.21 (d, *J* = 9.5 Hz), 108.38 (d, *J* = 25.1 Hz), 103.14 (s, 2C), 100.28, 60.21, 56.18 (s, 2C); HRMS-ESI (m/z): [M+H]^+^ calcd for C_18_H_18_N_2_O_3_F - 329.1301; found 329.1287; T_r_ = 3.32 min, purity >95%.

**7-fluoro-*N*-(3,4,5-trimethoxyphenyl)quinolin-4-amine (43)** prepared by method a to afford a light green solid (206 mg, 64%) ^1^H NMR (400 MHz, DMSO-*d*_6_) δ 11.17 (s, 1H), 8.97 (dd, *J* = 9.4, 5.6 Hz, 1H), 8.48 (d, *J* = 7.0 Hz, 1H), 7.91 (dd, *J* = 9.5, 2.6 Hz, 1H), 7.74 (ddd, *J* = 9.4, 8.4, 2.6 Hz, 1H), 6.84 (d, *J* = 7.0 Hz, 1H), 6.83 (s, 2H), 3.80 (s, 6H), 3.72 (s, 3H); ^13^C NMR (101 MHz, DMSO-*d*_6_) δ 165.47, 162.95, 155.09, 153.64, 143.26, 140.07, 139.95, 136.64, 132.65, 127.47 (d, *J* = 10.6 Hz), 116.48 (d, *J* = 24.4 Hz), 114.19, 105.09 (d, *J* = 25.1 Hz), 103.33(s, 2C), 100.41, 60.17, 56.15 (s, 2C); HRMS-ESI (m/z): [M+H]^+^ calcd for C_18_H_18_N_2_O_3_F - 329.1301; found 329.1289; T_r_ = 3.14 min

**5,7-difluoro-*N*-(3,4,5-trimethoxyphenyl)quinolin-4-amine (44)** prepared by method B to afford a bright yellow solid (191 mg, 55%) ^1^H NMR (400 MHz, DMSO-*d*_6_) δ 9.27 (s, 1H), 8.38 (d, *J* = 5.9 Hz, 1H), 7.61 (d, *J* = 9.5 Hz, 1H), 7.58 – 7.50 (m, 1H), 6.74 (s, 3H), 3.74 (s, 6H), 3.66 (s, 3H); ^13^C NMR (101 MHz, DMSO-*d*_6_) δ 163.53 (**d**, *J* = 15.8 Hz), 161.04 (d, *J* = 15.4 Hz), 158.49 (d, *J* = 14.6 Hz), 153.52, 151.53 – 150.55 (m), 149.16 – 147.34 (m), 146.11 (d, *J* = 12.6 Hz), 135.84, 134.09, 106.21 (d, *J* = 9.5 Hz), 105.38 (dd, *J* = 22.0, 4.1 Hz), 102.87, 102.62 – 101.01 (m, 1C), 60.19, 56.08 (s, 2C); HRMS-ESI (m/z): [M+H]^+^ calcd for C_18_H_17_F_2_N_2_O_3_ - 347.1207; found 347.1195; T_r_ = 3.19 min

**6-(*tert*-butyl)-*N*-(3,4,5-trimethoxyphenyl)quinolin-4-amine (45)** prepared by method B to afford a grey solid (224 mg, 67%) ^1^H NMR (400 MHz, DMSO-*d*_6_) δ 8.66 (s, 1H), 8.42 (d, *J* = 6.6 Hz, 1H), 8.10 – 7.98 (m, 2H), 6.82 (d, *J* = 3.0 Hz, 3H), 3.80 (s, 6H), 3.72 (s, 3H), 1.43 (s, 9H); ^13^C NMR (101 MHz, DMSO-*d*_6_) δ 153.91, 153.59 (s, 2C), 149.58, 143.22, 136.12, 133.72, 131.35, 121.49, 118.94, 117.15, 103.09 (**s**, 2C), 100.50, 60.22, 56.14 (s, 2C), 35.42, 31.15 (**s**, 3C); HRMS-ESI (m/z): [M+H]^+^ calcd for C_22_H_26_N_2_O_3_ - 367.2022; found 367.2009; T_r_ = 4.59 min, purity >95%.

**4-((3,4,5-trimethoxyphenyl)amino)quinoline-6-carbonitrile (46)** prepared by method A to afford a yellow solid (245 mg, 69%) ^1^H NMR (400 MHz, DMSO-*d*_6_) δ 11.38 (s, 1H), 9.49 (d, *J* = 1.3 Hz, 1H), 8.55 (d, *J* = 7.1 Hz, 1H), 8.33 (dd, *J* = 8.8, 1.5 Hz, 1H), 8.24 (d, *J* = 8.8 Hz, 1H), 6.98 (d, *J* = 7.1 Hz, 1H), 6.83 (s, 2H), 3.80 (s, 6H), 3.73 (s, 3H); ^13^C NMR (101 MHz, DMSO-*d*_6_) δ 154.88, 153.66 (s, 2C), 144.40, 140.38, 136.72, 134.65, 132.38, 130.58, 121.89, 117.92, 116.85, 108.95, 102.96 (s, 2C), 101.75, 60.18, 56.16 (s, 2C); HRMS-ESI (m/z): [M+H]^+^ calcd for C_19_H_18_N_3_O_3_ - 336.1348; found 336.1334; T_r_ = 3.11 min, purity >95%.

**6-(methylsulfonyl)-*N*-(3,4,5-trimethoxyphenyl)quinolin-4-amine (47)** prepared by method B to afford a dark yellow solid (231 mg, 72%) ^1^H NMR (400 MHz, DMSO-*d*_6_) δ 11.38 (s, 1H), 9.54 (s, 1H), 8.55 (d, *J* = 6.2 Hz, 1H), 8.47 – 8.16 (m, 2H), 6.99 (d, *J* = 6.1 Hz, 1H), 6.83 (s, 2H), 3.76 (d, *J* = 31.3 Hz, 9H), 3.44 (s, 3H); ^13^C NMR (101 MHz, DMSO-*d*_6_) δ 155.34, 154.02 (s, 2C), 145.80, 142.36, 138.73, 136.79, 133.43 (s, 2C), 130.02, 125.19, 117.41, 103.14 (s, 2C),102.25, 60.61, 56.56 (s, 2C), 44.00; HRMS-ESI (m/z): [M+H]^+^ calcd for C_19_H_21_N_2_O_5_S - 389.1171; found 389.1156; T_r_ = 2.98 min, purity >95%.

**6-methoxy-*N*-(3,4,5-trimethoxyphenyl)quinolin-4-amine (48)** prepared by method A to afford a light green solid (313 mg, 89%) ^1^H NMR (400 MHz, DMSO-*d*_6_) δ 10.75 (s, 1H), 8.40 (d, *J* = 6.9 Hz, 1H), 8.19 (d, *J* = 2.6 Hz, 1H), 8.03 (d, *J* = 9.2 Hz, 1H), 7.67 (dd, *J* = 9.2, 2.6 Hz, 1H), 6.86 (d, *J* = 6.9 Hz, 1H), 6.82 (s, 2H), 3.99 (s, 3H), 3.81 (s, 3H), 3.73 (s, 3H); ^13^C NMR (101 MHz, DMSO-*d*_6_) δ 157.94, 154.03, 153.65 (s, 2C), 140.69, 136.44, 133.03(s, 2C), 125.30, 122.06, 118.28 (s, 2C), 103.28, 102.68, 100.00, 60.19, 56.48, 56.15 (s, 2C); HRMS-ESI (m/z): [M+H]^+^ calcd for C_19_H_20_N_2_O_4_ - 340.1423; found 341.1486; T_r_ = 3.55 min, purity >95%.

**6,7-dimethoxy-*N*-(3,4,5-trimethoxyphenyl)quinolin-4-amine (49)** prepared by method A to afford a light green solid (285 mg, 86%) ^1^H NMR (400 MHz, DMSO-*d*_6_) δ 10.56 (s, 1H), 8.32 (d, *J* = 6.9 Hz, 1H), 8.11 (s, 1H), 7.44 (s, 1H), 6.92 – 6.70 (m, 3H), 3.98 (d, *J* = 8.9 Hz, 6H), 3.80 (s, 6H), 3.72 (s, 3H); ^13^C NMR (101 MHz, DMSO-*d*_6_) δ 154.51, 153.61 (s, 2C), 153.29, 149.36, 140.01, 136.29, 135.44, 133.19 (s, 2C), 111.45 (s, 2C), 103.26 (s, 2C), 99.59, 60.19, 56.66, 56.16, 56.14 (s, 2C); HRMS-ESI (m/z): [M+H]^+^ calcd for C_20_H_23_N_2_O_5_ - 371.1607; found 371.1589; T_r_ = 3.60 min, purity >95%.

**7-methoxy-*N*-(3,4,5-trimethoxyphenyl)quinolin-4-amine (50)** prepared by method A to afford an off white solid (278 mg, 79%) ^1^H NMR (400 MHz, DMSO-*d*_6_) δ 10.81 (s, 1H), 8.71 (d, *J* = 9.4 Hz, 1H), 8.38 (d, *J* = 7.0 Hz, 1H), 7.46 (d, *J* = 2.5 Hz, 1H), 7.41 (dd, *J* = 9.3, 2.5 Hz, 1H), 6.80 (s, 2H), 6.76 (d, *J* = 7.0 Hz, 1H), 3.96 (s, 3H), 3.80 (s, 6H), 3.72 (s, 3H); ^13^C NMR (101 MHz, DMSO-*d*_6_) δ 163.36, 155.09, 154.02 (s, 2C), 141.13, 136.88, 133.36 (s, 2C), 125.88, 118.54, 111.73, 103.75 (s, 2C), 100.48, 100.01, 60.61, 56.57 (s, 2C), 56.44; HRMS-ESI (m/z): [M+H]^+^ calcd for C_19_H_20_N_2_O_4_ - 340.1423; found 341.1485; T_r_ = 3.41 min, purity >95%.

**7-(trifluoromethyl)-*N*-(3,4,5-trimethoxyphenyl)quinolin-4-amine (51)** prepared by method A to afford a bright yellow solid (239 mg, 73%) ^1^H NMR (400 MHz, DMSO-*d*_6_) δ 11.12 (s, 1H), 8.99 (d, *J* = 8.9 Hz, 1H), 8.62 (d, *J* = 7.0 Hz, 1H), 8.45 (s, 1H), 8.14 (dd, *J* = 8.9, 1.5 Hz, 1H), 6.99 (d, *J* = 7.0 Hz, 1H), 6.83 (s, 2H), 3.81 (s, 6H), 3.73 (s, 3H); ^13^C NMR (101 MHz, DMSO-*d*_6_) δ 154.85, 153.71 (s, 2C), 144.39, 138.32 (d, *J* = 31.7 Hz, CF_3_), 136.71, 132.55, 125.80, 124.56, 122.29 (d, *J* = 3.5 Hz), 119.31, 118.25, 105.45, 103.09 (s, 2C), 101.68, 60.19, 56.17 (s, 2C); HRMS-ESI (m/z): [M+H]^+^ calcd for C_19_H_18_F_3_N_2_O_3_ - 379.1270; found 379.1253; T_r_ = 3.81 min, purity >95%.

**4-((3,4,5-trimethoxyphenyl)amino)quinoline-7-carbonitrile (52)** prepared by method A to afford a yellow solid (224 mg, 63%) ^1^H NMR (400 MHz, DMSO-*d*_6_) δ 11.49 (s, 1H), 9.55 (s, 1H), 8.54 (d, *J* = 7.1 Hz, 1H), 8.32 (dd, *J* = 8.8, 1.5 Hz, 1H), 8.25 (d, *J* = 8.8 Hz, 1H), 6.97 (d, *J* = 7.1 Hz, 1H), 6.84 (s, 2H), 3.80 (s, 6H), 3.72 (s, 3H); ^13^C NMR (101 MHz, DMSO-*d*_6_) δ 154.94, 153.63 (s, 2C), 144.19, 140.25, 136.71, 134.67, 132.37, 130.70, 121.72, 117.89, 116.82, 108.93, 102.99 (s, 2C), 101.73, 60.17, 56.15 (s, 2C); HRMS-ESI (m/z): [M+H]^+^ calcd for C_19_H_18_N_3_O_3_ - 336.1348; found 336.1333; T_r_ = 3.04 min, purity >95%.

***N*-(3-bromophenyl)-6,7-dimethoxyquinolin-4-amine (53)** prepared by method B to afford a colourless solid (183 mg, 57%) ^1^H NMR (400 MHz, DMSO-*d*_6_) δ 9.06 (s, 1H), 8.31 (d, *J* = 5.3 Hz, 1H), 7.69 (s, 1H), 7.50 (t, *J* = 1.9 Hz, 1H), 7.35 (ddd, *J* = 8.1, 2.1, 1.1 Hz, 1H), 7.28 (t, *J* = 7.9 Hz, 1H), 7.23 (s, 1H), 7.19 (ddd, *J* = 7.8, 1.9, 1.1 Hz, 1H), 6.91 (d, *J* = 5.3 Hz, 1H), 3.89 (d, *J* = 15.2 Hz, 6H); ^13^C NMR (101 MHz, DMSO-*d*_6_) δ 151.78, 148.35, 148.10, 145.89, 145.40, 143.25, 131.09, 125.05, 123.39, 122.00, 119.58, 114.55, 108.07, 102.15, 101.35, 56.02, 55.54; HRMS-ESI (m/z): [M+H]^+^ calcd for C_17_H_16_N_2_O_2_Br - 359.0395; found 359.0380; T_r_ = 3.99 min, purity >95%.

**6,7-dimethoxy-*N*-(3-methoxyphenyl)quinolin-4-amine (54)** prepared by method B to afford a colourless solid (180 mg, 65%) ^1^H NMR (400 MHz, DMSO-*d*_6_) δ 10.23 (s, 1H), 8.31 (d, *J* = 6.4 Hz, 1H), 8.08 (s, 1H), 7.43 (s, 1H), 7.40 (t, *J* = 8.4 Hz, 1H), 7.06 – 6.98 (m, 2H), 6.87 (ddd, *J* = 8.4, 2.4, 0.9 Hz, 1H), 6.81 (d, *J* = 6.4 Hz, 1H), 3.99 (s, 3H), 3.94 (s, 3H), 3.79 (s, 3H); ^13^C NMR (101 MHz, DMSO-*d*_6_) δ 160.21, 153.60, 150.89, 148.97, 142.28, 139.89, 138.59, 130.37, 116.22, 112.48, 111.31, 109.90, 102.44, 102.38, 100.01, 56.55, 55.92, 55.27; HRMS-ESI (m/z): [M+H]^+^ calcd for C_18_H_19_N_2_O_3_ - 311.1396; found 311.1382; T_r_ = 3.65 min, purity >95%.

***N*-(3-ethynylphenyl)-6,7-dimethoxyquinolin-4-amine (55)** prepared by method A to afford a dark colourless solid (193 mg, 71%) ^1^H NMR (400 MHz, DMSO-*d*_6_) δ 10.94 (s, 1H), 8.35 (d, *J* = 6.9 Hz, 1H), 8.25 (s, 1H), 7.60 (q, *J* = 1.4 Hz, 1H), 7.58 – 7.52 (m, 2H), 7.51 – 7.43 (m, 2H), 6.76 (d, *J* = 6.9 Hz, 1H), 4.34 (s, 1H), 3.99 (d, *J* = 19.4 Hz, 6H); ^13^C NMR (101 MHz, DMSO-*d*_6_) δ 154.58, 152.85, 149.41, 139.76, 138.07, 135.29, 130.21, 129.97, 128.14, 125.88, 123.15, 111.85, 102.98, 99.73, 99.31, 82.60, 81.87, 56.85, 56.14; HRMS-ESI (m/z): [M+H]^+^ calcd for C_19_H_17_N_2_O_2_ - 305.1290; found 305.1278; T_r_ = 3.75 min, purity >95%.

**1-(3-((6,7-dimethoxyquinolin-4-yl)amino)phenyl)ethan-1-one (56)** prepared by method A to afford a colourless solid (193 mg, 67%) HRMS-ESI (m/z): [M+H]^+^ calcd for C_19_H_19_N_2_O_3_ - 323.1396

**6-7-dimethoxy-*N*-(3,4,5-trimethoxyphenyl)quinolin-4-amine (57)** prepared by method C to afford a bright yellow solid (261mg, 79%) ^1^H NMR (400 MHz, DMSO-*d*_6_) δ 11.41 (s, 1H), 8.81 (s, 1H), 8.35 (s, 1H), 7.37 (s, 1H), 7.09 (s, 2H), 4.00 (d, *J* = 13.2 Hz, 6H), 3.79 (s, 6H), 3.70 (s, 3H); ^13^C NMR (101 MHz, DMSO-*d*_6_) δ 158.15, 156.17, 152.67 (s, 3C), 150.12, 148.59, 135.89, 132.65, 107.16, 104.04, 103.06 (s, 2C), 99.79, 60.17, 57.02, 56.43, 56.07 (s, 2C); HRMS-ESI (m/z): [M+H]^+^ calcd for C_19_H_22_N_3_O_5_ - 372.1559; found 372.1542; T_r_ = 3.26 min, purity >95%.

### Differential Scanning Fluorimetry (DSF) Assays

Starting from a 100 μM stock, GAK (14-351) was diluted to 1 μM in buffer 100 mM K_2_HPO_4_ pH7.5 containing 150 mM NaCl, 10% glycerol and 1X dye (Applied Biosystems catalog 4461806). The protein/dye mixture was transferred to a 384-well PCR microplate with 20 μL per well. Compounds at 10 μM in DMSO were added next, in 20 nL volume, using a liquid handling device setup with a pin head to make a final 10 μM compound in the assay plate. The final DMSO concentration in all wells was 0.1%, including the reference well with DMSO only.

Thermal shift data was measured in a qPCR instrument (Applied Biosystems QuantStudio 6) programmed to equilibrate the plate at 25 °C for 5 minutes followed by ramping the temperature to 95 °C at a rate of 0.05 °C per second. Data was processed on Protein Thermal shift software (Applied Biosystems) fitting experimental curves to a Boltzmann function to calculate differential thermal shifts (ΔT_m_) referenced to protein/dye in 0.1% DMSO only. The PKIS was screened against a purified GAK construct according to previously reported procedures (Fedorov, Niesen et al. 2012).

### Ligand binding displacement Assays

Inhibitor binding was determined using a binding-displacement assay, which tests the ability of the inhibitors to displace a fluorescent tracer compound from the ATP binding site of the kinase domain. Inhibitors were dissolved in DMSO and dispensed as 16-point, 2x serial dilutions in duplicate into black multi-well plates (Greiner). Each well contained either 0.5 nM or 1 nM biotinylated kinase domain protein ligated to streptavidin-Tb-cryptate (Cisbio), 12.5 nM or 25 nM Kinase Tracer 236 (ThermoFisher Scientific), 10 mM Hepes pH 7.5, 150 mM NaCl, 2 mM DTT, 0.01% BSA, 0.01% Tween-20. Final assay volume for each data point was 5 μL, and final DMSO concentration was 1%. The plate was incubated at room temperature for 1.5 hours and then read using a TR-FRET protocol on a PheraStarFS plate reader (BMG Labtech). The data was normalized to 0% and 100% inhibition control values and fitted to a four parameter dose-response binding curve in GraphPad Software (version 7, La Jolla, CA, USA). The determined IC_50_ values were converted to *K*_i_, values using the Cheng-Prusoff equation and the concentration and *K_d_* values for the tracer (previously determined).

### Competition binding assays

Competition binding assays were performed at DiscoverX as described previously (Davis, Hunt et al. 2011). Kinases were produced either as fusions to T7 phage3, or were expressed as fusions to NF-κB in HEK-293 cells and subsequently tagged with DNA for PCR detection. In general, full-length constructs were used for small, single-domain kinases, and catalytic domain constructs including appropriate flanking sequences were used for multidomain kinases. Briefly, for the binding assays, streptavidin-coated magnetic beads were treated with biotinylated affinity ligands to generate affinity resins. The liganded beads were blocked to reduce non-specific binding and washed to remove unbound ligand.

Binding reactions were assembled by combining kinase, liganded affinity beads, and test compounds prepared as 100× stocks in DMSO. DMSO was added to control assays lacking a test compound. Assay plates were incubated at 25 °C with shaking for 1 h, and the affinity beads were washed extensively to remove unbound protein. Bound kinase was eluted in the presence of nonbiotinylated affinity ligands for 30 min at 25 °C with shaking. The kinase concentration in the eluates was measured by quantitative PCR. The KINOMEscan^®^ panel of approximately 400 wild type human kinase assays was run as a single measurement at 1 μM. K_d_ values were determined using 11 serial threefold dilutions of test compound and a DMSO control.

### Molecular Modelling

Molecular modelling was performed using Schrödinger Maestro software package (Small-Molecule Drug Discovery Suite 2016-3, Schrödinger, LLC, New York, NY, 2016.) Structures of small molecules were prepared using and the LigPrep module of Schrodinger suite employing OPLS3 force for all computations. X-ray crystal structure for the GAK (pdb:4Y8D) was pre-processed using the protein preparation wizard of Schrödinger suite in order to optimize the hydrogen bonding network (Kovackova, Chang et al. 2015).

Prior to Glide docking, the grid box was centered using corresponding x-ray ligand as template. The ligand docking was performed using default SP settings of Schrodinger Glide with additional hydrogen bond constraints to NH of CYS126 (hinge residue). Graphical illustrations were generated using MOE software (Molecular operating environment (MOE), 2016.8; chemical computing group Inc., 1010 Sherbooke St. West, Suite #910, Montreal, QC, Canada, H3A 2R7, 2016.), Maestro (WaterMap) or PyMOL (The PyMOL Molecular Graphics System, Version 1.8 Schrödinger, LLC).

### Hydration Site Analysis

Hydration site analysis calculated with WaterMap (Schrödinger Release 2016-3: WaterMap, Schrödinger, LLC, New York, NY, 2016.). The 4Y8D structure was prepared with Protein Preparation Wizard (as above). Waters were analyzed within 5 Å of the co-crystallized ligand, and the 2 ns simulation was conducted with OPLS3 force field.

### Gas Phase/OPLS3 force field modelling

Torsional barriers of compounds of over N-C linger were estimated using molecular mechanics calculations employing Rapid Torsion Scan functionality of Maestro suite with angle increment of 15 degrees with OPLS3 force field. QM optimized gas phase geometries for compounds shown in table 2 were calculated with Jaguar software. Initial geometries were based on the rapid torsion scan and full geometry optimization was calculated using B3LYP level of theory and 6-31G** basis set with standard geometry optimization setting of Jaguar (Schrödinger Release 2016-3: Jaguar, Schrödinger, LLC, New York, NY, 2016).

## SUPPLEMENTARY INFORMATION

### ACKNOWLEDGMENT

The SGC is a registered charity (number 1097737) that receives funds from AbbVie, Bayer Pharma AG, Boehringer Ingelheim, Canada Foundation for Innovation, Eshelman Institute for Innovation, Genome Canada, Innovative Medicines Initiative (EU/EFPIA), Janssen, Merck & Co., Novartis Pharma AG, Ontario Ministry of Economic Development and Innovation, Pfizer, São Paulo Research Foundation-FAPESP, Takeda, and Wellcome Trust. David H. Drewry is gratefully acknowledged for stimulating discussions. We thank the EPSRC UK National Crystallography Service for funding and the collection of the crystallographic data. We would also like to thank the Biocenter Finland/DDCB for financial support towards the goals of our work and the CSC-IT Center for Science Ltd. (Finland) for the allocation of computational resources. In addition, we also thank Dr. Brandie M. Ehrmann for LC-MS/HRMS support provided by the Mass Spectrometry Core Laboratory at the University of North Carolina at Chapel Hill.

## REFERENCES

Abel, R., T. Young, R. Farid, B. J. Berne and R. A. Friesner (2008). “Role of the active-site solvent in the thermodynamics of factor Xa ligand binding.” J Am Chem Soc 130(9): 2817–2831.

Abrami, L., S. Liu, P. Cosson, S. H. Leppla and F. G. van der Goot (2003). “Anthrax toxin triggers endocytosis of its receptor via a lipid raft-mediated clathrin-dependent process.” J Cell Biol 160(3): 321–328.

Aleksandrowicz, P., A. Marzi, N. Biedenkopf, N. Beimforde, S. Becker, T. Hoenen, H. Feldmann and H. J. Schnittler (2011). “Ebola virus enters host cells by macropinocytosis and clathrin-mediated endocytosis.” J Infect Dis 204 Suppl 3: S957–967.

Barbieri, E., P. P. Di Fiore and S. Sigismund (2016). “Endocytic control of signaling at the plasma membrane.” Curr Opin Cell Biol 39: 21–27.

Bekerman, E. and S. Einav (2015). “Infectious disease. Combating emerging viral threats.” Science 348(6232): 282–283.

Bellei, B., A. Pitisci, E. Migliano, G. Cardinali and M. Picardo (2014). “Pyridinyl imidazole compounds interfere with melanosomes sorting through the inhibition of cyclin G-associated Kinase, a regulator of cathepsins maturation.” Cell Signal 26(4): 716–723.

Bridges, A. J. (1999). “The rationale and strategy used to develop a series of highly potent, irreversible, inhibitors of the epidermal growth factor receptor family of tyrosine kinases.” Curr Med Chem 6(9): 825–843.

Cohen, P. and D. R. Alessi (2013). “Kinase drug discovery‐‐what’s next in the field?” ACS Chem Biol 8(1): 96–104.

Davis, M. I., J. P. Hunt, S. Herrgard, P. Ciceri, L. M. Wodicka, G. Pallares, M. Hocker, D. K. Treiber and P. P. Zarrinkar (2011). “Comprehensive analysis of kinase inhibitor selectivity.” Nat Biotechnol 29(11): 1046–1051.

Dolomanov, O. V., L. J. Bourhis, R. J. Gildea, J. A. K. Howard and H. Puschmann (2009). “Olex2: A complete structure solution, refinement and analysis program.” J. Appl. Cryst. 42: 339–341.

Drewry, D. H., C. I. Wells, D. Andrews, Richard Angell, H. Al-Ali, A. D. Axtman, S. J. Capuzzi, J. Elkins, P. Ettmayer, M. Frederiksen, O. Gileadi, N. Grey, A. Hooper, S. Knapp, S. Laufer, U. Luecking, S. Muller-Knapp, E. Muratov, D. Rajiah, K. Saikatendu, D. Trieber, W. J. Zuercher and T. M. Willson (2017). “Progress Towards a Public Chemogenomic Set for Protein Kinases: A Call for Contributions.” bioaRxiv.

Dumitriu, A., J. C. Latourelle, T. C. Hadzi, N. Pankratz, D. Garza, J. P. Miller, J. M. Vance, T. Foroud, T. G. Beach and R. H. Myers (2012). “Gene expression profiles in Parkinson disease prefrontal cortex implicate FOXO1 and genes under its transcriptional regulation.” PLOS Genet 8(6): e1002794.

Dumitriu, A., C. D. Pacheco, J. B. Wilk, K. E. Strathearn, J. C. Latourelle, S. Goldwurm, G. Pezzoli, J. C. Rochet, S. Lindquist and R. H. Myers (2011). “Cyclin-G-associated kinase modifies alpha-synuclein expression levels and toxicity in Parkinson’s disease: results from the GenePD Study.” Hum Mol Genet 20(8): 1478–1487.

Dzamko, N., J. Zhou, Y. Huang and G. M. Halliday (2014). “Parkinson’s disease-implicated kinases in the brain; insights into disease pathogenesis.” Front Mol Neurosci 7: 57.

Fabian, M. A., W. H. Biggs, 3rd, D. K. Treiber, C. E. Atteridge, M. D. Azimioara, M. G. Benedetti, T. A. Carter, P. Ciceri, P. T. Edeen, M. Floyd, J. M. Ford, M. Galvin, J. L. Gerlach, R. M. Grotzfeld, S. Herrgard, D. E. Insko, M. A. Insko, A. G. Lai, J. M. Lelias, S. A. Mehta, Z. V. Milanov, A. M. Velasco, L. M. Wodicka, H. K. Patel, P. P. Zarrinkar and D. J. Lockhart (2005). “A small molecule-kinase interaction map for clinical kinase inhibitors.” Nat Biotechnol 23(3): 329–336.

Fedorov, O., F. H. Niesen and S. Knapp (2012). “Kinase inhibitor selectivity profiling using differential scanning fluorimetry.” Methods Mol Biol 795: 109–118.

Greener, T., X. Zhao, H. Nojima, E. Eisenberg and L. E. Greene (2000). “Role of cyclin G-associated kinase in uncoating clathrin-coated vesicles from non-neuronal cells.” J Biol Chem 275(2): 1365–1370.

Haile, P. A., B. J. Votta, R. W. Marquis, M. J. Bury, J. F. Mehlmann, R. Singhaus, Jr., A. K. Charnley, A.S. Lakdawala, M. A. Convery, D. B. Lipshutz, B. M. Desai, B. Swift, C. A. Capriotti, S. B. Berger, M. K. Mahajan, M. A. Reilly, E. J. Rivera, H. H. Sun, R. Nagilla, A. M. Beal, J. N. Finger, M. N. Cook, B. W. King, M. T. Ouellette, R. D. Totoritis, M. Pierdomenico, A. Negroni, L. Stronati, S. Cucchiara, B. Ziolkowski, A. Vossenkamper, T. T. MacDonald, P. J. Gough, J. Bertin and L. N. Casillas (2016). “The Identification and Pharmacological Characterization of 6-(tert-Butylsulfonyl)-N-(5-fluoro-1H-indazol-3-yl)quinolin-4-amine (GSK583), a Highly Potent and Selective Inhibitor of RIP2 Kinase.” J Med Chem 59(10): 4867–4880.

Kanaoka, Y., S. H. Kimura, I. Okazaki, M. Ikeda and H. Nojima (1997). “GAK: a cyclin G associated kinase contains a tensin/auxilin-like domain.” FEBS Lett 402(1): 73–80.

Kimura, S. H., H. Tsuruga, N. Yabuta, Y. Endo and H. Nojima (1997). “Structure, expression, and chromosomal localization of human GAK.” Genomics 44(2): 179–187.

Kooistra, A. J., G. K. Kanev, O. P. van Linden, R. Leurs, I. J. de Esch and C. de Graaf (2016). “KLIFS: a structural kinase-ligand interaction database.” Nucleic Acids Res 44(D1): D365–371.

Korolchuk, V. I. and G. Banting (2002). “CK2 and GAK/auxilin2 are major protein kinases in clathrin-coated vesicles.” Traffic 3(6): 428–439.

Kovackova, S., L. Chang, E. Bekerman, G. Neveu, R. Barouch-Bentov, A. Chaikuad, C. Heroven, M. Sala, S. De Jonghe, S. Knapp, S. Einav and P. Herdewijn (2015). “Selective Inhibitors of Cyclin G Associated Kinase (GAK) as Anti-Hepatitis C Agents.” J Med Chem 58(8): 3393–3410.

Louie, J. and J. F. Hartwig (1995). “Palladium-Catalyzed Synthesis of Arylamines from Aryl Halides. Mechanistic Studies Lead to Coupling in the Absence of Tin Reagents.” Tetrahedron Lett. 36(21): 3609–3612.

Manning, G., D. B. Whyte, R. Martinez, T. Hunter and S. Sudarsanam (2002). “The protein kinase complement of the human genome.” Science 298(5600): 1912–1934.

Mercer, J., M. Schelhaas and A. Helenius (2010). “Virus entry by endocytosis.” Annu Rev Biochem 79: 803–833.

Neveu, G., A. Ziv-Av, R. Barouch-Bentov, E. Berkerman, J. Mulholland and S. Einav (2015). “AP-2- associated protein kinase 1 and cyclin G-associated kinase regulate hepatitis C virus entry and are potential drug targets.” J Virol 89(8): 4387–4404.

Palatinus, L. and G. Chapuis (2007). “Superflip — A computer program for solution of crystal structures from x-ray diffraction data in arbitrary dimension.” J. Appl. Cryst. 40: 786–790.

Papatheodorou, P., C. Zamboglou, S. Genisyuerek, G. Guttenberg and K. Aktories (2010). “Clostridial glucosylating toxins enter cells via clathrin-mediated endocytosis.” PLOS One 5(5): e10673.

Ravez, S., O. Castillo-Aguilera, P. Depreux and L. Goossens (2015). “Quinazoline derivatives as anticancer drugs: a patent review (2011 - present).” Expert Qpin Ther Pat 25(7): 789–804.

Ray, M. R., L. A. Wafa, H. Cheng, R. Snoek, L. Fazli, M. Gleave and P. S. Rennie (2006). “Cyclin G-associated kinase: a novel androgen receptor-interacting transcriptional coactivator that is overexpressed in hormone refractory prostate cancer.” Int J Cancer 118(5): 1108–1119.

Rhodes, S. L., J. S. Sinsheimer, Y. Bordelon, J. M. Bronstein and B. Ritz (2011). “Replication of GWAS associations for GAK and MAPT in Parkinson’s disease.” Ann Hum Genet 75(2): 195–200.

Rudolf, A. F., T. Skovgaard, S. Knapp, L. J. Jensen and J. Berthelsen (2014). “A comparison of protein kinases inhibitor screening methods using both enzymatic activity and binding affinity determination.” PLOS One 9(6): e98800.

Sakurai, M. A., Y. Ozaki, D. Okuzaki, Y. Naito, T. Sasakura, A. Okamoto, H. Tabara, T. Inoue, M. Hagiyama, A. Ito, N. Yabuta and H. Nojima (2014). “Gefitinib and luteolin cause growth arrest of human prostate cancer PC-3 cells via inhibition of cyclin G-associated kinase and induction of miR-630.” PLOS One 9(6): e100124.

Sheldrick, G. M. (2015). “Crystal structure refinement with SHELXL.” Acta Crystallogr C Struct Chem 71(Pt 1): 3–8.

Sheldrick, G. M. (2015). “SHELXT - integrated space-group and crystal-structure determination.” Acta Crystallogr A Found Adv 71(Pt 1): 3–8.

Shimizu, H., I. Nagamori, N. Yabuta and H. Nojima (2009). “GAK, a regulator of clathrin-mediated membrane traffic, also controls centrosome integrity and chromosome congression.” J Cell Sci 122(Pt 17): 3145–3152.

Sikka, V., V. K. Chattu, R. K. Popli, S. C. Galwankar, D. Kelkar, S. G. Sawicki, S. P. Stawicki and T. J. Papadimos (2016). “The Emergence of Zika Virus as a Global Health Security Threat: A Review and a Consensus Statement of the INDUSEM Joint working Group (JWG).” J Glob Infect Dis 8(1): 3–15.

Sloan, R. D., B. D. Kuhl, T. Mesplede, J. Munch, D. A. Donahue and M. A. Wainberg (2013). “Productive entry of HIV-1 during cell-to-cell transmission via dynamin-dependent endocytosis.” J Virol 87(14): 8110–8123.

Solomon, V. R. and H. Lee (2011). “Quinoline as a privileged scaffold in cancer drug discovery.” Curr Med Chem 18(10): 1488–1508.

Sorrell, F. J., M. Szklarz, K. R. Abdul Azeez, J. M. Elkins and S. Knapp (2016). “Family-wide Structural Analysis of Human Numb-Associated Protein Kinases.” Structure 24(3): 401–411.

Susa, M., E. Choy, X. Liu, J. Schwab, F. J. Hornicek, H. Mankin and Z. Duan (2010). “Cyclin G-associated kinase is necessary for osteosarcoma cell proliferation and receptor trafficking.” Mol Cancer Ther 9(12): 3342–3350.

Tabara, H., Y. Naito, A. Ito, A. Katsuma, M. A. Sakurai, S. Ohno, H. Shimizu, N. Yabuta and H. Nojima (2011). “Neonatal lethality in knockout mice expressing the kinase-dead form of the gefitinib target GAK is caused by pulmonary dysfunction.” PLOS One 6(10): e26034.

Takada, Y. and K. Matsuo (2012). “Gefitinib, but not erlotinib, is a possible inducer of Fra-1-mediated interstitial lung disease.” Keio J Med 61(4): 120–127.

van Linden, O. P., A. J. Kooistra, R. Leurs, I. J. de Esch and C. de Graaf (2014). “KLIFS: a knowledge-based structural database to navigate kinase-ligand interaction space.” J Med Chem 57(2): 249–277.

Wu, P., T. E. Nielsen and M. H. Clausen (2015). “FDA-approved small-molecule kinase inhibitors.” Trends Pharmacol Sci 36(7): 422–439.

Young, T., R. Abel, B. Kim, B. J. Berne and R. A. Friesner (2007). “Motifs for molecular recognition exploiting hydrophobic enclosure in protein-ligand binding.” Proc Natl Acad Sci U S A 104(3): 808–813.

